# Near-complete genomes for nine haplochromine cichlid fishes reveal a novel centromeric satellite structure organised around a pair of inverted elements

**DOI:** 10.64898/2026.06.30.735501

**Authors:** Pio Sierra, Chenxi Zhou, Bettina Fischer, Sing Wei Lim, Moritz Blumer, Maxon Ngochera, Richard Durbin

## Abstract

The haplochromine cichlid fishes of Lake Malawi form one of the most dramatic examples of recent rapid radiation in vertebrates. Here we describe nine new diploid telomere-to-telomere (T2T) genome sequences generated using ultra-long ONT reads, which include 78 ungapped chromosomes. We provide accurate annotations of transposable elements and tandem repeats, identify rDNA cluster regions and putative centromeres, and confirm previously reported large chromosomal inversions. The putative centromeres are primarily composed of satellite tandem arrays of previously reported 237 bp repeats, but notably on most chromosomes these are organised in a novel structure in which four blocks of satellites in alternating orientation are separated by an inverted pair of ∼15 kb sequences we term “centroids”, which have similarity to a non-autonomous DNA transposable element and containing potential CENP-B binding boxes. The methylation dip region indicating the likely active centromere always lies between the centroids, whose separation is almost always around 200 kb (interquartile range 151-221kb). A structurally equivalent but non-homologous organisation is seen in the distantly related *Etroplus* cichlid genera from South Asia. By comparing these structures across chromosomes and species, we suggest how they may have evolved, and potentially how they could contribute to the rampant sympatric speciation seen in these species, based on meiotic drive and chromosome missegregation.

## Introduction

Cichlid fishes are a family (Cichlidae) of teleosts distributed widely in Africa and South America, with presence also in Madagascar and the Indian subcontinent. The cichlids of the East African great lakes, in particular Lake Malawi, have long been considered one of the most striking examples of recent adaptive radiation (Malinsky et al., 2018). With more than 500 endemic species, Lake Malawi has the largest diversity of fish species of any lake in the world (Lyons et al., 2015). They display extensive phenotypic diversity despite base-level sequence divergence as low as 0.1%–0.25% between species pairs (Malinsky et al., 2018; Vernaz et al., 2021). This has led to suggestions that larger scale structural variation, in the form of multiple haplotypes which include insertion and absence polymorphisms, may have played an important role in adaptive variation (McGee et al., 2020). There are several molecular mechanisms that can have contributed to creating such variation in the genome. One of them is the varied transposable element (TE) content of haplochromine cichlids, which had been suggested already to be an important driver of key regulatory changes, facilitating rapid phenotypic change and possibly speciation (Brawand et al., 2014). An initial comparison of eight draft genome assemblies suggested that up to 10% of the genome is structurally variable between Malawi cichlids, dominated by TE insertion polymorphism (Quah et al., 2025). However the detailed study of the effect of structural variants in this radiation has been hindered by the lack of highly contiguous reference genomes that can help us elucidate the complex diversity present in these species. In particular several segregating large chromosomal inversions that have recently been reported within the radiation (Blumer et al., 2025) were not apparent in the draft assemblies, nor do they support detailed analysis of the evolutionary dynamics of rapidly evolving repeat sequences including the centromeres.

Here we present new, high quality, diploid reference genomes for eight species of Lake Malawi cichlids: *Astatotilapia calliptera, Aulonocara stuartgranti, Diplotaxodon limnothrissa, Labeotropheus fuelleborni, Labeotropheus trewavasae, Lethrinops chilingali, Maylandia zebra* and *Rhamphochromis chilingali,* plus *Astatotilapia burtoni* from Lake Tanganyika and adjacent rivers as an outgroup. For the first time this provides access to full centromere-spanning sequences for Lake Malawi cichlids, and we find in them not only the expected diversity of these rapidly evolving regions of the genome, but also a conserved structure, to our knowledge not previously reported in animal or plant centromeres. We discuss in detail the composition and conservation of this organisation, and provide a model of how it may have evolved and the potential implications for the evolution of this radiation.

### New genome assemblies for nine species of Lake Malawi cichlids

Using deep long and ultra-long sequencing data (57-107x coverage) we assembled diploid genomes for nine samples of different haplochromine species, in each case providing a primary sequence with maximal contiguity and a full-length secondary sequence (Supplementary Table 1; see methods for details). We used Hi-C data from the same individual to scaffold the *Aulonocara stuartgranti* (Astu) assembly to chromosomes, orienting and naming them according to established convention (Malinsky et al., 2018), and this together with full chromosome sequences (see below) to complete chromosomal assignment for the other species.

The final result for the nine primary assemblies includes 78 telomere-to-telomere (T2T) chromosomes that are ungapped with telomeric sequence at both ends, 60 more chromosomes with no gaps but missing one or both telomeres, and 79 gaps in the remaining 56 chromosomes (40 in *Astatotilapia calliptera* and 14 in *Labeotropheus fuelleborni*), with only two gaps in *Lethrinops chilingali* (Figure 1a). All 22 chromosomes have a T2T instance in at least one species, ranging from one for chromosomes 1, 3 and 17 up to seven for chr11. In addition there are two more T2T chromosomes (2 and 6) in the secondary assembly for *Lethrinops chilingali*.

**Figure 1:**
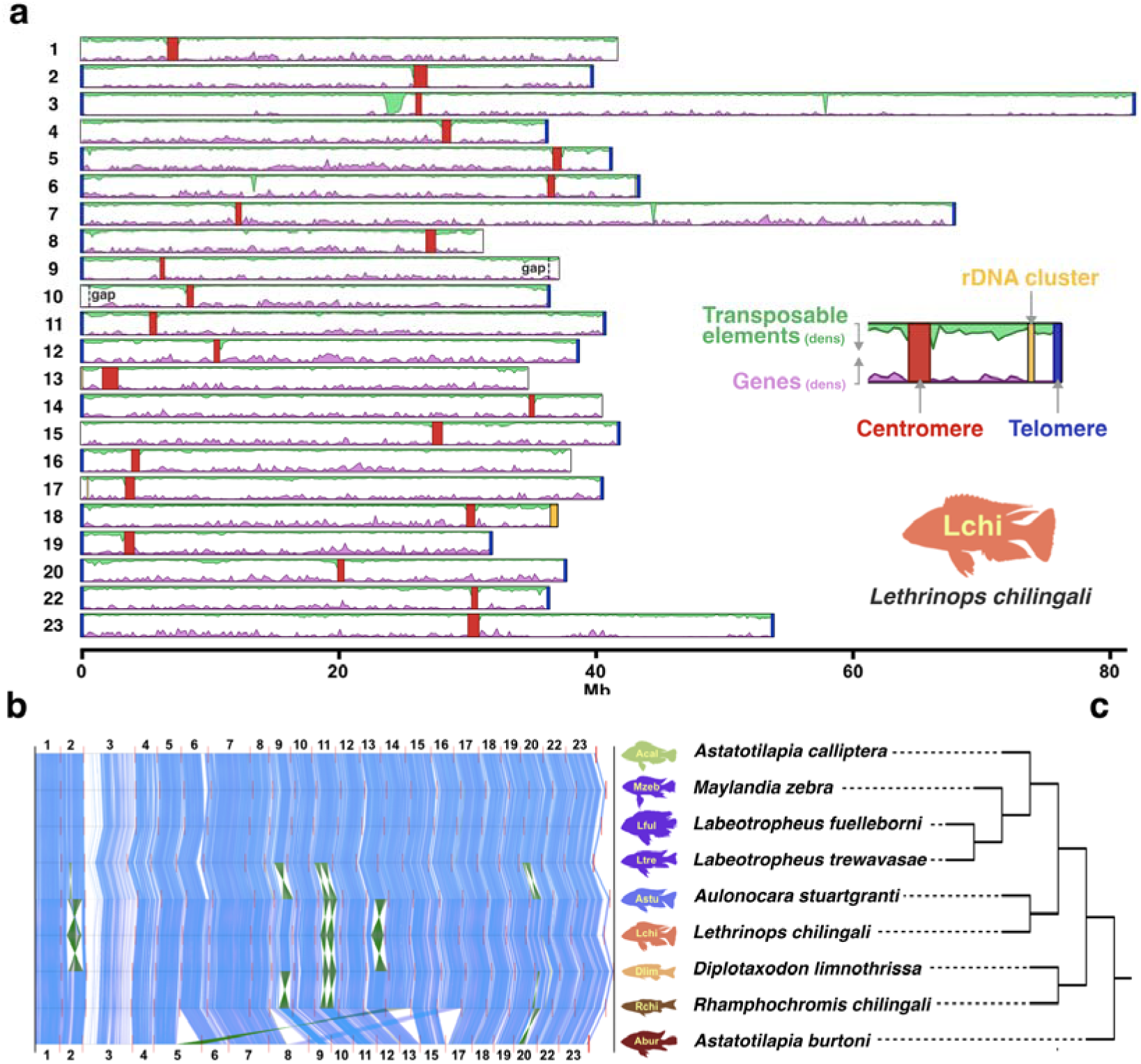
Nine new assemblies of haplochromine cichlids. **a,** Karyotype plot of *Lethrinops chilingali*, showing density of gene content (lower histogram) and transposable element content (upper histogram, inverted), centromeres, telomeres and the locations of the two gaps (chr9 and chr10). The red rectangles show the location of each centromere. **b,** Synteny plots between the nine genomes produced with Mumemto (Shivakumar and Langmead, 2025). Inversions appear as hourglasses. White spaces belong to highly repetitive regions where no shared unique matches could be found. *Astatotilapia burtoni* has chromosome 14 fused with chromosome 5 and chromosome 16 fused with chromosome 8. In both cases the lower chromosome number has been kept. **c,** Species cladogram of the nine species in **b**, obtained with Roadies (Gupta et al., 2025). We confirmed several segregating large chromosomal inversions that had been reported in some species of the radiation (Blumer et al., 2024) (Figure 1b), and the fusions of chromosomes 5-14 and 8-16 in *Astatotilapia burtoni* (Roberts et al., 2016).

We annotated transposable elements (TEs) in the new genomes using a combined TE library of 2,738 families obtained from all genomes (see Methods). We found no significant differences in the total amount of TE content between the species, but we observed that there are some TE families which are particular only of a certain species, while others are shared between all of them. In addition we have also annotated tandem repeats, including the telomeric repeat. The vertebrate telomeric repeat TTAGGG is present at 254 (5’: 124, 3’:130) out of 392 ends of chromosomal scaffolds. There was no chromosome end for which the telomeric repeat was missing in all assemblies. We found rDNA arrays on the short arms of chromosomes 6, 13, 17 and 18 in most species, and occasionally on other acrocentric chromosomes (Supplementary Table 2). The sizes varied, but it is possible that the assemblies were collapsed due to identity so further investigation would be required before drawing conclusions from the sizes.

No large translocations were found between chromosomes, and in general chromosome synteny is highly conserved between species, with the exceptions of the already mentioned inversions and and the two pairs of chromosome fusions previously reported in the more distant *Astatotilapia burtoni* (Conte et al., 2021) (Figure 1b).

### The structure of haplochromine centromeres

We identified putative centromeric regions in all but one of the 196 primary assembly chromosomes, based on them containing regions dominated by large tandem satellite arrays of a 237 bp repeat, which has 84% identity across 223 bp to a previously reported 238 bp centromeric repeat unit found in related Tilapia species (Franck et al., 1992) (see Methods for details). The 237 bp monomer (mwcisat-237) has an average CG content of 37.6%, slightly under the 41% average of the genome.

Of the 195 chromosomal centromeric regions, 167 were assembled without gaps (Supplementary Table 3). The one missing centromere, on chromosome 6 of *Labeotropheus fuelleborni* was found in the unplaced scaffolds for that assembly. These putative centromeres were in consistent syntenic positions across all species, except for one copy being lost in each of the two fused chromosomes in *Astatotilapia burtoni* (discussed further below). Based on the ratio between the length of both chromosome arms r = q/p we identified 3 metacentric chromosomes (r < 1.5), 3 submetacentric chromosomes (1.5 < r < 3), and 16 acrocentric chromosomes (r > 3) (Figure 1a), compatible with previous karyotypes from a number of cichlids, including some from the lakes of East Africa (Poletto et al., 2010). According to their results the majority of species have 22 chromosomes, most of them acrocentric, with a small number of metacentric or submetacentric chromosomes. Chromosome 3, the largest chromosome in these species, includes a large 2 Mb tandem array of partial LINE elements upstream of the centromere. All pericentromeric regions are enriched for recent TE insertions (Figure 1a). Centromere sizes, measured as the length occupied by the tandem repeat (TR) arrays of the 237 base monomer, range from less than 0.5 Mb up to 3 Mb, with the largest being found in chromosome 10 of *Astatotilapia burtoni*.

### Two inverted elements define the structure of these centromeres

Tandem repeats (TRs) of the mwcisat-237 are not the only components of haplochromine centromeres. In addition to a few occasional random TE insertions in some of them, we found a particular sequence, which from now on we call a “centroid”, that appears twice in most centromeres, in inverted relative orientation, flanked by tandem arrays of the centromeric satellite (Figure 2a). 98 out of 196 centromeres include two centroids, 43 have one centroid, and 55 have none (Supplementary Table 4). No two species had exactly the same distribution of centroids in their centromeres (Supplementary Table 4), but there were strongly shared patterns. There are two chromosomes, 3 and 9, for which both centroids were present in all nine primary assemblies, and three more, 1, 17 and 23, where only one centroid was missing, in all three cases in the assembly of *Astatotilapia calliptera*, which has lower contiguity. In contrast, centroids are almost completely absent in chr 4 (only one copy in *Lethrinops chilingali*), chr 13 (a full pair in *Astatotilapia burtoni*), and in chr 18 (a full pair in *Aulonocara stuartgranti* and 1 copy in *Maylandia zebra*).

**Figure 2:**
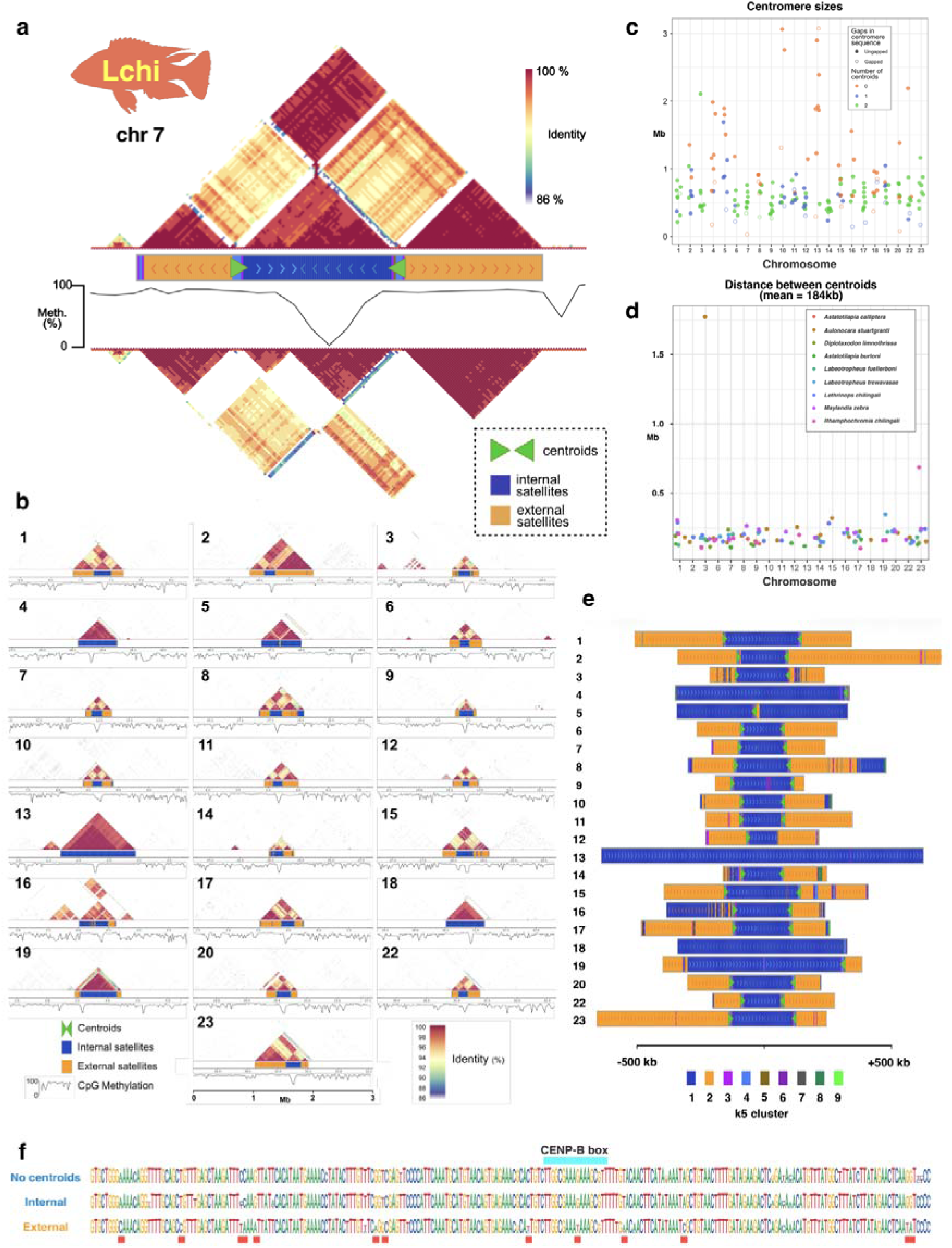
Centromeres of *Lethrinops chilingali*. **a,** Dot plot visualization of the centromere on chromosome 7 of *Lethrinops chilingali* obtained with ModDotPlot (Sweeten et al., 2024). The colors on the upper and lower triangles indicate the identity between different regions of the centromere. The lower, inverted ModDotPlot uses a variant in which the reverse complement is not considered for matching, which highlights how the tandem repeats are split into four blocks of alternating orientation, as indicated in the central band. The central band also shows the positions and orientations of the centroids (green arrows), and colours the tandem repeat regions as in **c**. The central histogram shows the level of CpG methylation in sliding 20kb windows, with the Centromere Dip Region (CDR) situated between the centroids. **b,** Dotplots, kmer composition of the satellites, centroid locations, and CpG methylation for all 22 chromosomes of *Lethrinops chilingali*. **c,** Distribution of centromere sizes by chromosome across the nine species in Figure 1. The colors indicate the presence of none, one or two centroids. Centromere sequences with gaps are labelled with an open rather than a filled circle. **d,** Distance between pairs of centroids (when present) on each centromere. The two outliers on chromosome 3 of *Aulonocara stuartgranti* and chromosome 23 of *Rhamphochromis chilingali* are due to large insertions present in those centromeres. **e,** Composition of the tandem repeats across all *L. chilingali* centromeres. Each centromere was divided in 2370 b windows (10x satellite size) and the 5-mer composition of each window was used to cluster them into 9 clusters. For all chromosomes with two centroids, the clusters 1 and 2 label the satellite sequences inside and outside the centroids respectively. Clusters 3 to 9 label TE insertions and the centroids. In the absence of centroids, the centromere tends to be composed only of the internal satellites (cluster 1). **f,** Consensus sequences of the inner (internal) and outer (external) satellite sequences of centromeres with two centroids of *Lethrinops chilingali* and the satellite sequences of centromeres without two centroids (No centroids) highlighting their putative CENP-B box (blue) and base positions which are more divergent between the internal and external motifs.

### Centroids provide structure to the centromeres

We identified the boundaries of the centromeres and the positions of the centroids in all 198 chromosomes. Strikingly, the centroids are flanked by divergent TR arrays of centromeric satellite sequence (Figure 2a). When both centroids are present, the sequence between the centroids is therefore divided into two TR arrays of convergent satellite sequences that meet at a point between them, not always at the same distance from each centroid (Supplementary Table 3). Looking outwards from the centroids, towards the pericentromeric regions, one or several TR arrays appear.

Not all centromeres contain two centroids: some have either one or none (Figure 2b; Supplementary Figures 8-29). The centromeres can therefore be classified into one of three types. First, centromeres where the two centroids are present, creating a symmetric pattern of satellites oriented in opposite directions as described above. These are almost all under 1Mb in size (Figure 2c), with the distance between centroids typically 100kb to 250kb, with mean 183kb and interquartile range 151-221kb (Figure 2d). Second, centromeres with no centroids, with a large dominating TR array in which all the satellites are in the same direction, more similar in sequence to the internal arrays of the two-centroid class. These are typically larger than the previous group, with the majority between 1Mb and 3Mb in size. Third, centromeres which have just one centroid, with most TR arrays composed of satellites in the same direction and of the internal type, but only in a few cases longer than 1Mb.

To compare the composition of the TRs we clustered windows of 10 satellite sequences (2370 bases) by their 5-mer composition. Surprisingly we found that the different segments cluster with each other on different chromosomes according to their relative position in the centromere (Figure 2e). Groups from the external TRs, when centroids are present, cluster together in most cases regardless of the chromosome. Groups from the internal TRs also cluster together regardless of the chromosome but, critically, they also cluster with the groups of satellites from centroidless centromeres, or those in which one centroid is missing (Figure 2f). We do not see consistent higher-order repeat structure locally within the TR arrays, but there are short patterns of low level divergence, contained within one of the separate blocks of satellite repeats flanking a centroid (Supplementary Figure 30).

To identify the putative location of the active centromere we studied the CpG methylation patterns to find the Centromeric Dip Regions (CDRs), regions where the centromeric satellites, which are typically heavily methylated, experience a strong reduction in methylation (Altemose et al., 2022; Gershman et al., 2022). In chromosomes with two centroids, the CDR is always present in the region between them (Figure 2b), although it is not necessarily at the point of convergence of the two internal TR arrays. This suggests that the centroids could play some role in determining active centromere position. When the centroids are missing, the CDR is present in a large, highly homogeneous TR array in which there is no inversion of the satellites, with all of them oriented in the same sense.

### Sequence composition of the centroids

We created a consensus of the centroid sequences and identified several features within it (Figure 3a). We found no significant Open Reading Frames (ORFs) in the centroid, and mapping of RNAseq data could not confirm any transcription from it. However, it shows various structural features suggesting a relationship to DNA transposable elements. Starting from the outside of the centromeres, the consensus sequence of the centroid is composed of several copies, each more divergent than the previous one, of the satellite repeat. Then comes in most centroids, but not all, a sequence that is repeated later in opposite direction, forming an inverted repeat, which includes between the inverted sequences a composite of almost two and a half satellite-like sequences (Figures 3a,b). Another feature that can be identified is a region with homology to a PiggyBac transposase (DDE_Tnp_1_7, Pfam domain PF13843), but very degraded so no longer constituting a functional ORF. Towards the inside end, the centroid starts to include the same satellite sequences that will compose the rest of the centromere, but they are separated from the tandem array itself by a short stretch of non-satellite sequence. Of these satellite-like sequences, the most proximal from the centromere internal part (at position 12,343 in the consensus centroid) includes a CENP-B box which is shared by the satellites. The centroids also include two G homopolymers of 12 and 15 bases that could result in G-quadruplexes or i-motifs (Matos-Rodrigues et al., 2023). In addition to these features, the centroids are rich in small palindromes, besides the large inverted repeat, that could form cruciform structures.

**Figure 3:**
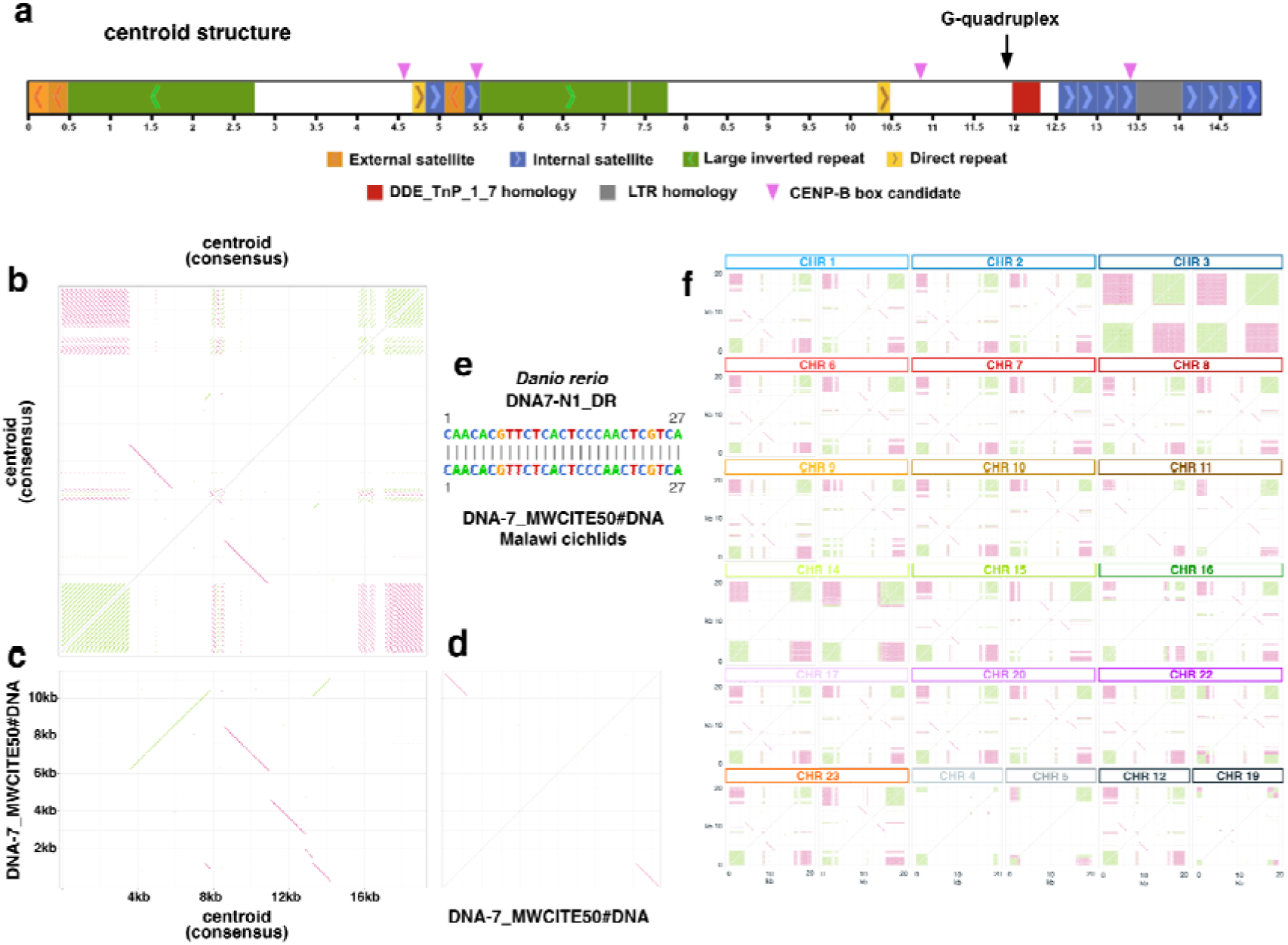
Structure and composition of centroids. **a,** Structural features identified on the consensus sequence obtained from the alignment of all centroids in the nine genomes. The centroid is flanked by tandem repeats of satellites (internal and external) some of which may be part of the centroid. **b,** Self dotplot of the consensus centroid sequence. **c** Dotplot comparing the sequences of the consensus centroid and a non autonomous DNA TE (DNA-7_MWCITE50) identified in cichlids of Lake Malawi. **d,** Dotplot of the non autonomous transposable element DNA-7_MWCITE50. **e,** Identity between the first 27 bases of DNA-7_MWCITE50 and non-autonomous DNA element DNA7-N1_DR identified in *Danio rerio*. **f,** Self dotplots of all 36 centroids in the primary assembly of fLetChi2. The last four centroids at the bottom don’t have a complementary centroid in their centromere.

We compared the centroid sequence to the library of TEs we created for Lake Malawi cichlids and we found long stretches of near-identical (94%) sequence similarity between the centroid sequence and a non-autonomous transposable element (DNA-7_MWCITE50#DNA) (Figure 3c,d). This element is classified by RepeatClassifier as a member of the Tigger superfamily, but the insertions present in the genome include a 7 to 8 basepair target site duplication (TSD), instead of the canonical 2bp TA TSD for the Tigger superfamily, suggesting this element might contain remnants of a Tigger TE, but has been mobilized recently by another type of TE, potentially Zisupton, as the termini of the elements matched the motifs identified in termini for elements of this superfamily, conserved across fishes to zebrafish (Figure 3e) (Böhne et al., 2012; Storer et al., 2021).

The centroids vary across chromosomes, with each centroid more similar to the other one in the same chromosome than to those in other chromosomes (Figure 3f). Some of them include the full sequence described in the consensus, while others appear to either be just part of it, or have been degraded. Chromosome 3 has two centroids, but these are particularly degraded, missing the first half that includes the inverted repeat. The centroids in chromosomes 11 and 14 lack the core of cryptic satellites inside the inverted repeat, which is missing completely in chr 14. As mentioned, the presence of the two centroids is an indication of conservation of a certain structure of the centromere, with a large expansion of a single TR array when one or both of them are missing. Interestingly, the most glaring exception to the conservation of synteny between pairs of centroids happens in *Lethrinops chilingali* in chromosome 16 where the homology between them is less conserved. In this case the satellites also do not follow the general rule and satellites that cluster better with the internal ones can be found already in the 5’ external TR array.

Centroids also tend to be conserved between homologous chromosomes of different species (Figure 4). Indeed, in general sequence similarity is more conserved between species, as expected of transmission through vertical descent than between the pair on a chromosome; this is exemplified by chromosome 2, chromosome 3 except in *Rhamphochromis chilingali*, 6 except in *Aulonocara stuartgranti*. Nevertheless, in some cases, there is more similarity between paired centroids of one chromosome. This is most frequent for *Astatotilapia burtoni*, which is a divergent outgroup with respect to the other species all from Lake Malawi, although we also see this occasionally even where there is a closely related species, such as on chromosome 16 of *Labeotropheus fuelleborni*. These observations suggest that there is an intermittent process of homogenisation via gene conversion between the two centroids within a chromosome.

**Figure 4:**
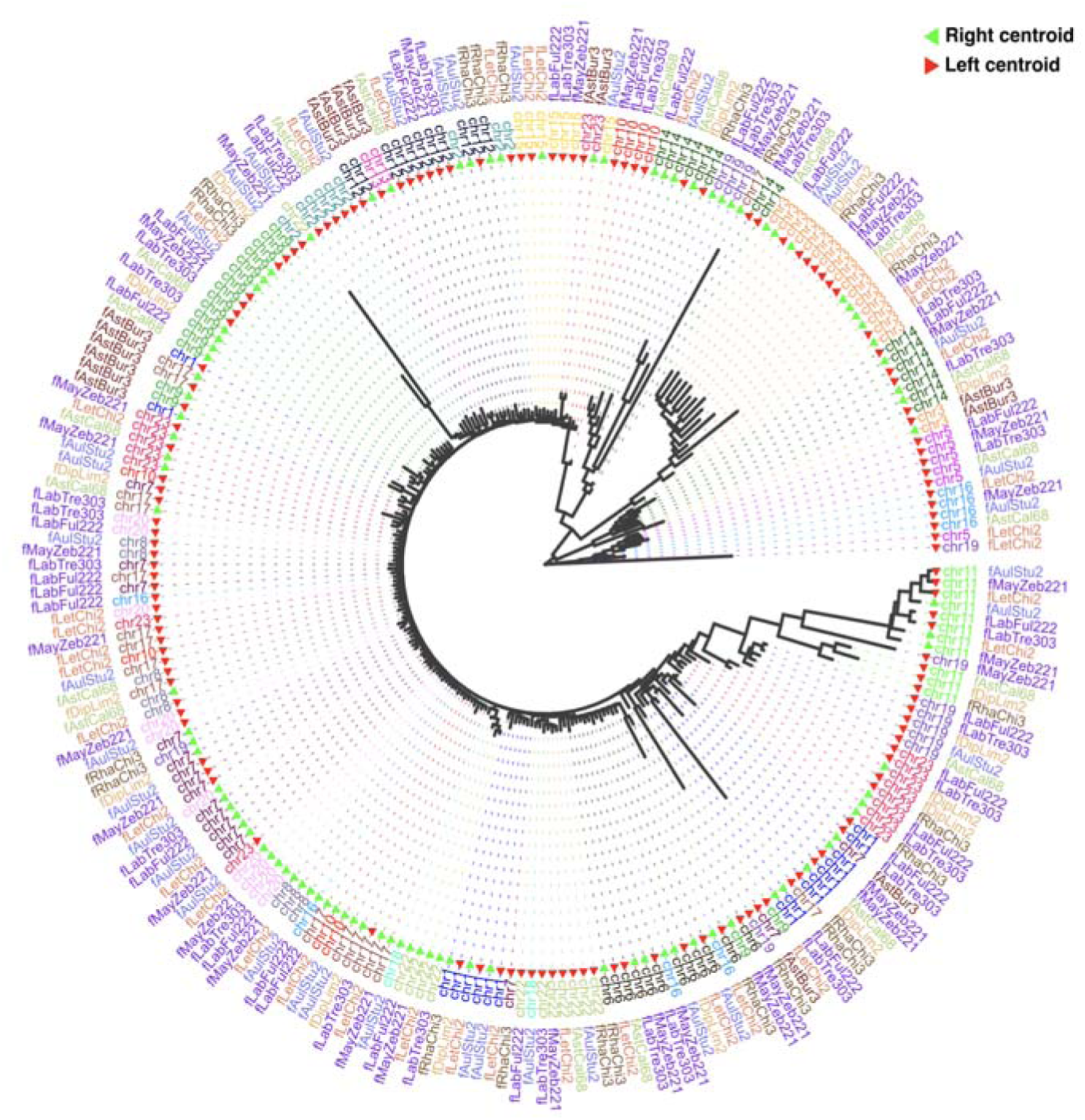
Centroid relatedness dendrogram. Dendrogram of the 317 centroid sequences found in the primary haplotypes of the nine cichlid genomes in Figure 1. From outside to inside: Genome name, chromosome name, relative position of the centroid.

### One centromere has been lost in each of the chromosome fusions in *Astatotilapia burtoni*

The karyotype of *Astatotilapia burtoni* differs from the Lake Malawi species in that there are two fusions of chromosomes. Most African cichlids have a diploid chromosome number of 2n=44, this suggests that the diploid chromosome number of 2n=40 of *A. burtoni* is the derived state (Conte et al., 2021), with ancestral chromosomes 5 and 14 fusing to form chromosome 5 in *A. burtoni* and ancestral chromosomes 8 and 16 fusing to form chromosome 8 in *A. burtoni*. The 5-14 fusion was found to possibly include an XY sex determination locus in laboratory populations (Böhne et al., 2016; Roberts et al., 2016), but those results were not reproduced in wild populations (Lichilín et al., 2023).

In our new *A. burtoni* genome we obtained ungapped sequences for both fusions, enabling us to study their evolutionary trajectory. We confirmed that both cases result from fusions between the short arms of the ancestral chromosomes. To evaluate the loss or duplication of genome sequence we compared a region of 12 Mb surrounding the putative joining area with a composite of two 6Mb regions from the ends of the short arms of the homologous chromosomes of *Aulonocara stuartgranti* (Figure 5a,b). We found that the two cases are similar. First, the whole chromosomes have fused, close to their telomeres, not at the acrocentric centromeres. Second, the actual telomeric sequences found in the non-fused chromosomes are missing in the fusions, together with a small part of the subtelomeric sequence. Third, in both cases one of the centromeres from the unfused chromosomes (original chromosomes 14 and 16) is in the same exact place as in the fused chromosome, while the other centromere (original chromosomes 5 and 8) is essentially missing. We say “essentially” because in both cases a few copies of the centromeric satellites remain, less than 5kb total, strongly suggesting that the old centromeres were there and have been removed, rather than that the fusion happened before these centromeres appeared. This is consistent with removal of one of the DNA centromeric sequences being necessary to stabilize dicentric chromosomes (Tyler-Smith et al., 1993).

**Figure 5:**
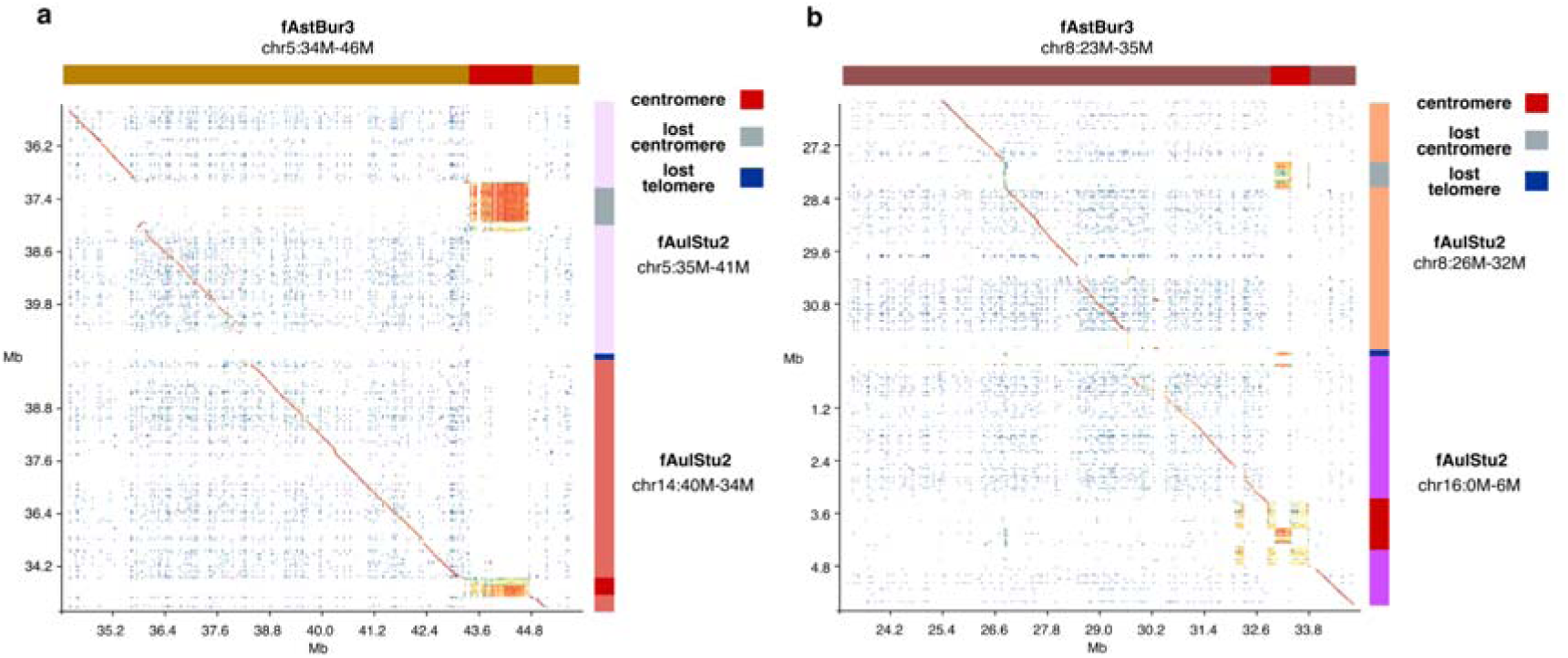
Centromeres in *Astatotilapia burtoni* chromosome fusions. **a,** Dotplot showing sequence identity between 12Mb around the chromosome fusion of *Astatotilapia burtoni* chromosome 5 and the concatenated sequences of the homologous chromosome ends in chromosome 14 and chromosome 5 in *Aulonocara stuartgranti*. The sequence of chromosome 14 is inverted. Red blocks indicate the location of the putative centromeres in those chromosomes. Blue blocks indicate the location of the telomere repeats in *Aulonocara stuartgranti.* The telomere repeats and one of the centromeres are missing in *Astatotilapia burtoni.* **b,** Same as **a** but for the fusion of chromosomes 8 and 16.

Compared to most Malawi species, *A. burtoni* is missing many centroids, including on chromosomes 5 and 8 (only 19 in total are present, 16 in pairs on 8 chromosomes and 3 singletons). The centromere of *A. burtoni* chromosome 5 (homologous to that chromosome 14 in other species) is not different from other centromeres that have lost their centroids. In contrast, the centromere for chromosome 8 (homologous to chromosome 16 in other species), has evolved to be composed of a satellite of 238 bases, which has an additional T base, outside the putative CENP-B box. All the copies of the satellite repeat on *A. burtoni* chromosome 8 have this larger length.

### Centroids with different sequence composition are found in other cichlids

To understand better the origin of these centromeres and their centroids we examined available public genomes for other cichlids. We identified putative centromeres in high quality assemblies for 12 other species (Figure 6a). We compared them according to three factors: a) Identity of their satellites compared to the 237 satellite in the Lake Malawi cichlids. b) Presence of centroids, defined as pairs of inverted non-satellite sequences flanking one or several highly identical tandem repeats of satellites that are shared between chromosomes. c) Orientation of the satellites as spreading from the centroids, with the region between the centroids consequently divided into two tandem repeats in opposite directions.

**Figure 6:**
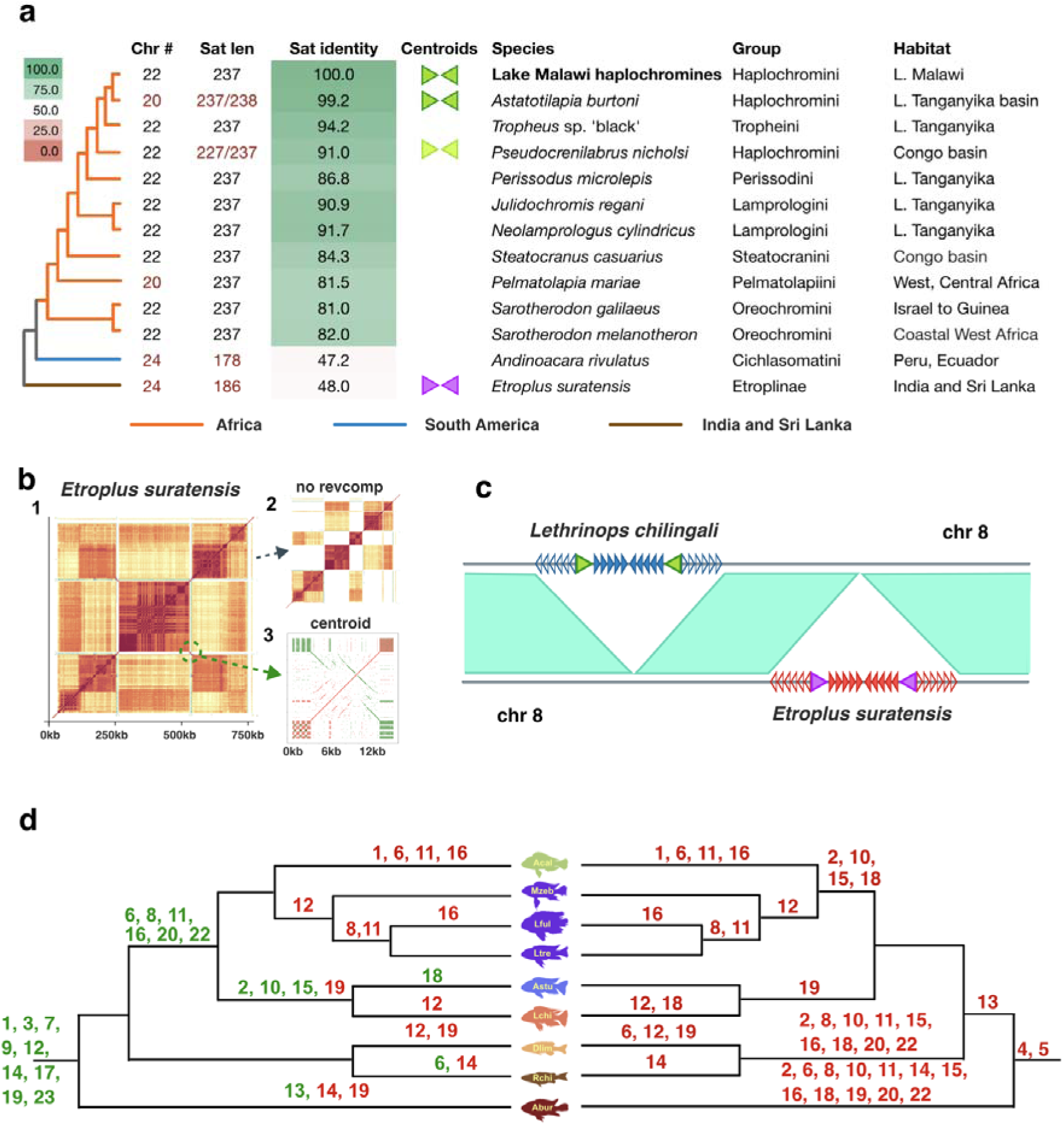
Centromeres and centroids in cichlid fishes. **a,** Species in which centroids and the 237bp satellite sequence are observed. The Tropheini tribe of Lake Tanganyika is nested within the tribe Haplochromini. The centroids found in *Etroplus suratensis* bear no homology to the centroids of Lake Malawi cichlids. **b,** *Etroplus suratensis*, a basal cichlid found in India and Sri Lanka, presents centromeres with the same structure of centroids and tandem repeats as the haplocromines: **1** ModDotPlot of chromosome X, **2** modified ModDotPlot without reverse-complement matches, revealing alternating orientation of satellite regions as in African species, **3** self-dotplot of the centroid. **c,** In spite of the shared structure, the satellite and the centroids have different sequences than their African counterparts, and the centromeres are also in a different position in the chromosomes. **d** Possible models of the evolution of centroids on Malawi cichlid chromosomes placed onto a consensus species phylogeny. To the left, a phylogenetic model in which gains (green) and losses (red) of the centroids occur; to the right, a model in which there are only losses.

We found the same centroid and tandem repeat structure in Lake Malawi haplochromine species sequenced by others, *Mchenga conophoros* and *Copadichromis virginalis*, but not *Tropheus* sp. *‘*black’ from the Tropheini tribe of Lake Tanganyika, which are within a different branch of the Haplochromini from the Malawi species, although their satellite sequence is nearly identical. However, in *Pseudocrenilabrus nicholsi*, an even more distantly related haplochromine, we found a single centromere on chromosome 9 that shares the structure of having a pair of inversely oriented centroids, though these are composed of the full DNA TE element that is scrambled in the centroids of Lake Malawi haplochromines. The centromeric satellites on this chromosome are essentially the same as those in the other haplochromines, while those on other chromosomes are 10bp shorter due to a single 10bp deletion.

Looking beyond Haplochromini, *Neolamprologus cylindricus* (Lamprologini) equally lacked any centroid or the inverted tandem repeat structures. In *Perissodus microlepis*, a more distant Perissodini from Lake Tanganika, we find insertions of the same DNA TE element in the centromeres, sometimes two in a centromere, but typically they are in the same orientation and, importantly, the tandem repeats between them are all in the same orientation, suggesting that they are just independent insertions in centromeres of a particular active or centrophilic element. We could not find any other candidates for centroids in other African cichlids, nor in *Andinoacara rivulatus*, a South American cichlid with a completely different 178 satellite repeat, and estimated age of divergence of 62.1 Ma (70.1–54.6 Ma) (Matschiner et al., 2020).

However, strikingly, *Etroplus suratensis*, a basal cichlid from India and Sri Lanka, which has a different satellite repeat of 186 bases, not homologous to the African or the American ones, with estimated age of divergence of 76.2 Ma (86.6–66.3 Ma) (Matschiner et al., 2020), has structurally very similar centromeres to the ones we identified in Malawi haplochromines (Figure 6b). They also are formed of a non-satellite sequence of approximately 15kb, paired in inverse orientation in each centromere, shared between centromeres, that divides the central part of the putative centromere into two satellite tandem repeat regions oriented in opposing directions. Remarkably, these centroids do not share any homology with the haplochromine centroids (Figure 6c), in this case being formed from LINE RTE transposable element insertions, among other features. Like the haplochromine cichlids, however, they do include a few copies of the satellites inside their sequence, flanked in both cases by inverted repeats.

The pattern of presence or absence of centroids on chromosomes within the species sequenced does not simply fall on the consensus phylogenetic tree relating the species (Figure 6d). A model in which there are both gains and losses (left side of Figure 6d) is more parsimonious than one involving just losses (right side), but still requires some repeated events. Models with only gains would require even more events and seem implausible.

Given the sequence sharing patterns in Figure 4, we note that the changes here, either gains or losses, are more likely to have occurred by long range gene conversion or introgression rather than de novo creation and convergent evolution; we know that hybridisation and introgression have occurred in the past in the Lake Malawi radiation (Malinsky et al., 2018).

## Discussion

In eukaryotes, centromeres are the chromosomal loci where kinetochores form to provide a connection between the DNA and the spindle microtubules during mitosis and meiosis. Centromeres are essential for cell division but in spite of this fact their sequences evolve rapidly and can vary drastically in their composition and organization, even between closely related species (Nagaki et al., 2025). The repetitive nature of most centromeres and their basic patterns of DNA composition have been known for more than 50 years (Kit, 1961), but it has not been until recently that the availability of high quality long and ultra-long reads has made it possible to accurately assemble the full sequences of centromeres (Naish et al., 2021; Altemose et al., 2022; Baalsrud et al., 2024; Chaudhry et al., 2024; Heuberger et al., 2024; Yoo et al., 2024; Xie et al., 2025).

In this work we have studied the DNA composition of assemblies with complete putative centromeric sequences from eight species of cichlids from Lake Malawi and a closely related outgroup. We have found in them a pattern of structural organisation that, to our knowledge, has not been previously reported. Our work so far has been based only on DNA sequencing data, and further analysis would be necessary to confirm the presence of CENP-A histones in the centromeres, though we can confirm the presence of a highly expressed CenpA-like gene. Still, we consider the clear signal of a hypomethylated region surrounded by hypermethylated satellite segments in all the putative centromeres, coupled with their chromosomal locations, consistent with what was previously reported in Lake Malawi cichlids karyotypes, as good indicators that they are likely to be the active centromeres of each chromosome.

The structure that we find, although not previously reported in animals, bears an intriguing resemblance to the small regional centromeres of fission yeast *Schizosaccharomyces pombe* (Figure 7a), in which an inverted pair of innermost repeats (imrs), each of size around 15kb, flank the active centromere marked by CENP-A, and themselves are flanked by outer repetitive sequence that is methylated to form heterochromatin (Xu et al., 2023). It has been suggested that this spatial inverted layout is necessary to properly fold the DNA loop, stabilizing the three-dimensional physical platform upon which the inner kinetochore complex is built (Brar and Amon, 2008). We propose that the paired centroids in cichlids (Figure 7b) could facilitate stable kinetochore localisation in a similar fashion. Intriguingly, Hoyt et al. have recently reported that LINE elements “demarcate neocentromere boundaries in humans, implicating transposable elements in restricting CENP-A domain spreading”; they do not report the orientation of these LINE elements (Hoyt et al., 2025).

**Figure 7:**
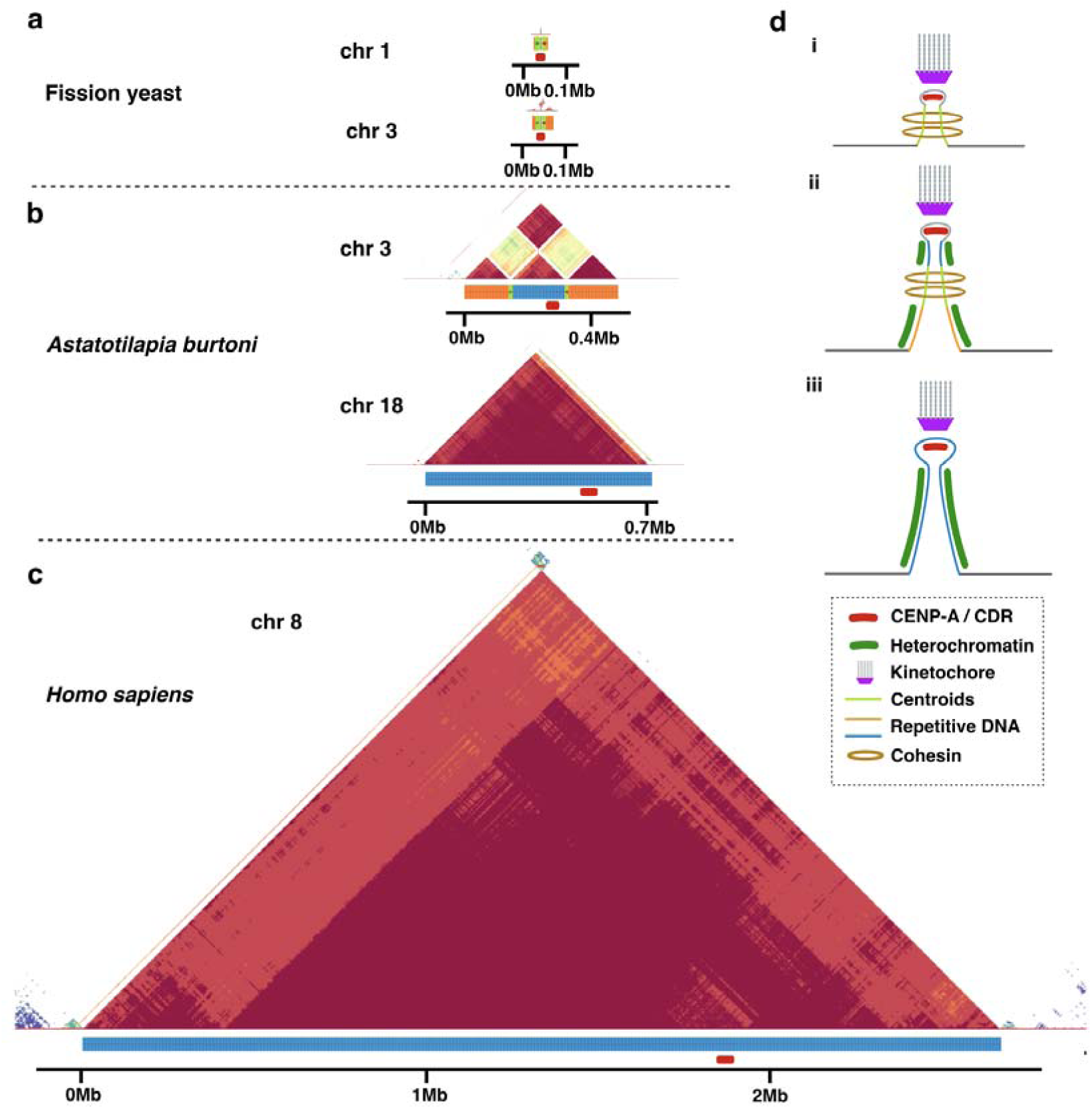
A model for the evolution of regional centromeres. **a,** Structure of two centromeres of fission yeast *Schizosaccharomyces pombe*. An inverted pair of innermost repeats (imr) flanks the central core (cc) of the regional centromeres. In chr 3 expansions of tandem repeats flank the innermost repeats. **b,** Two example centromeres from cichlids. **c,** Human chromosome 8 centromere. All of **a,b,c** are to the same scale. **d,** Alternative states for regional centromeres. **i,** The smallest regional centromeres are defined by a flanking pair of inverted repeats of around 15kb (imrs or centroids) that delimit the area of the active centromere in a larger region of heterochromatin. They help recruit Cenp-A and may provide a 3D structure that extrudes the active centromere, facilitating the pairing with the kinetochore and the orientation of the homologous chromosomes and sister chromatids during mitosis and meiosis. **ii,** The inclusion of satellite repeat sequences around the centroids is selected favorably to provide a base for constitutive heterochromatin that helps maintain the structure and isolates the active centromere. **iii,** eventually enough heterochromatin and the presence of Cenp-A are enough to maintain the centromere, and the centroids can be lost.

It does appear likely that the spacing of the inverted centroid pair is important, since that strongly tends to be in the range of 150-250kb, which we note is similar to the distribution of lengths of kinetochore sites in the human CHM1 and CHM13 cell lines inferred from CDRs and CENP-A binding (Logsdon et al., 2024), although our observed CDRs are typically smaller, around 50kb. Potentially selection on centroid spacing helps limit the size of the centromeric satellite arrays, which otherwise tend to expand by misaligned recombinational repair, at least in part via breakage-induced replication (Talbert and Henikoff, 2022; Showman et al., 2024) (Figure 7b,c). This results in larger arrays in the absence of paired centroids (Figure 2b,c). It is also striking that in the absence of centroids it is the inner satellite repeat that supports the active centromere which forms the major satellite array in Malawi cichlids (Figure 2e): the CDR is always within this repeat type (Figure 2b). We identified a potential CENP-B binding box in the centroids and inner satellite consensus, with a single base change in the outer satellite consensus (Figure 2f). Jiao et al. have recently reported functional conservation of CENP-B boxes in centromeric satellite sequence (Jiao et al., 2026).

The centroid organisation, while prevalent, is not always present in every chromosome, and hence is clearly not essential for centromere function. However, it appears to have arisen at least twice independently within cichlids, in East African haplochromines and in *Etroplus*, and potentially more than once independently in haplochromines given that the centroid observed in *Pseudocrenilabrus,* although related, is different from that found in Malawi cichlids. This raises the question of how this pattern arises. In all three instances, the centroid sequence appears to be derived from, or in the case of *Pseudocrenilabrus* the same as, that of a recently active transposable element. It is frequently observed that TEs insert into centromeric satellite regions (Naish et al., 2021; Wlodzimierz et al., 2023b; Courret et al., 2024), as indeed we see in *Perissodus*. Then selection on randomly inserted TEs could lead to fixation if they are functionally useful, as has been proposed in multiple other cases to underlie the establishment of a dispersed functional element (e.g. Ellison and Bachtrog, 2013; Carelli et al., 2022). It is tempting to suggest that the *Pseudocrenilabrus* case represents an early step in this process, since its sequence has not yet been selected away from the parental TE sequence to optimise centromeric function, and is not yet established on other chromosomes, although it has already reorganised satellite orientation. The fact that, although centroids seem to be mostly inherited vertically within their chromosomal location, they also occasionally convert within a chromosome or, more rarely, between chromosomes (Figure 4), providing a secondary mechanism for them to spread across chromosomes. Alongside these ways to establish centroids, the data suggest an ongoing pattern of centroid loss, falling back to a standard, predominantly unidirectional satellite-based centromere (Figure 6d).

Taken together, these observations suggest that a centroid-based centromeric sequence organisation may be functionally selected, but nevertheless a transient state in the dynamics of centromere evolution. Its discovery in, and high degree of variability within, the East African haplochromine cichlid radiation raises the question of whether it potentially actively contributes to adaptive speciation within this large species complex of many hundreds of species. It has been established that hybridisation occurs, or has occurred in the past, between many of these species (Joyce et al., 2011; Malinsky et al., 2018; Svardal et al., 2020), despite which they remain morphologically and ecologically distinct species. We observed that different species typically differ in which chromosomes have centroids. While it is plausible that the centroid state confers some form of advantage in mitosis where sister chromatids are paired, it also seems plausible that heterozygosity in centroid state, with one copy being centroid-based and the other having a single linear satellite cluster, might inhibit homologous pairing and hence be deleterious in meiosis, increasing the chance of missegregation; this would reduce hybrid fitness, supporting species separation. It would be possible to test this hypothesis by looking at the karyotypes of the sperm or unfertilised eggs of a laboratory hybrid fish, potentially by low coverage single cell genomic DNA sequencing. On a population genetic basis, this would also suggest that changes in centroid state are relatively hard to establish, leading to stronger differentiation between species in centroid state than at other loci. Of course, successful hybridisation and introgression events could transfer centroid state between species, as well as gene conversion as discussed above.

Finally, it is known that centromeres are highly dynamic at the sequence level, and indeed that it is possible to create and (more rarely) establish neocentromeres that move the whole position of the centromere on a chromosome (Scott and Sullivan, 2014; Kim, 2022). It is interesting to speculate about the potential role of centroid-like organisation in forming and stabilising such neocentromeres, perhaps originating from a fortuitously located pair of inverted TEs in heterochromatin (Figure 7d). Our results do not provide direct evidence for such a role - if we are correct that the centroids we found in *Pseudocrenilabrus* are new, then they inserted into an existing centromere location. However, we note that the discovery of the centroid-based centromeric sequence organisation was only made possible by the recent advent of high quality long-read based genome assemblies. These are now rapidly becoming available for thousands of species across the tree of life (Sequence locally, think globally: The Darwin Tree of Life Project | PNAS, 2022; Blaxter et al., 2025), including a rapidly increasing number that include full ungapped centromere sequences. We hypothesize that as these sequences are investigated in detail they may reveal additional examples of centroid-based centromeres, enabling further insight into the role that they can play in centromere evolution.

## Methods

### Library preparation and sequencing

For *Astatotilapia calliptera*, *Aulonocara stuartgranti*, *Rhamphochromis chilingali* and *Maylandia* sp. ‘pearly’ samples were obtained from aquarium stocks maintained at the University of Cambridge. *Labeotropheus fuelleborni* and *Labeotropheus trewavasae* samples were obtained from the wild in Lake Malawi, flash frozen and brought to the UK under an Access and Benefit sharing agreement between the University of Cambridge, the Government of Malawi, and the Wellcome Sanger Institute. A *Diplotaxodon limnothrissa* sample was obtained from the University of Hull aquarium. For *Diplotaxodon limnothrissa* and *Rhamphochromis chilingali* PacBio data were collected at the Wellcome Sanger Institute. For the others, extraction was performed as per Qiagen Genomic Tip 100/G protocol with minor modifications. Tissue was homogenised in a mortar with liquid Nitrogen, then subjected to lysis with RNAse for 1h at 37°C followed by Proteinase digestion for 2 h at 50°C. The lysate was then purified through the tips and all centrifuge steps were performed in microcentrifuge in 2mL aliquots. The quality and quantification of DNA was assessed with Nanodrop and Qubit BR DNA assay and size distribution with TapeStation Genomic Tape. High molecular weight (HMW) DNA was then repurified using the PacBio SRE kit (<25kb). The resulting HMW DNA was sequenced with PromethION.

### Genome assembly

For *Astatotilapia calliptera*, we ran HERRO (Stanojević et al., 2026) for ONT read error-correction. The error-corrected reads were assembled using Hifiasm (Cheng et al., 2021) with parameter ‘--primary’. Contigs representing haplotypic duplications were removed using purge_dups (Guan et al., 2020) with default parameters. Hi-C reads were mapped to the primary contigs using BWA-MEM2 (Vasimuddin et al., 2019) with parameters ‘-5 -S -P -C -p’ and processed using samtools (Li et al., 2009) ‘fixmate’ with parameters ‘-m -p -u’, then ‘sort’, and finally ‘markdup’ with parameters ‘-c’. The processed Hi-C alignments were used as input for YaHS (Zhou et al., 2023) for scaffolding with default parameters. The resulting genome assembly was then aligned to the published *Astatotilapia calliptera* reference genome (GCA_900246225.6) using FastGA (Myers et al., 2025) to fix scaffold order and orientation to ensure consistency in chromosome nomenclature. For the other species, we used Hifiasm in ONT mode (parameters ‘--primary --ont’) to assemble genomes directly from the ONT reads, without prior error-correction using HERRO. As Hi-C data were not available for these species, we mapped contigs to the *Astatotilapia calliptera* assembly to order contigs into chromosomes. Mitochondrial genomes were assembled using Oatk (Zhou et al., 2025) from error-corrected reads generated by either HERRO or Hifiasm. Nuclear genome assemblies were then aligned to the corresponding mitochondrial genomes to identify and remove mitochondrial contigs from the nuclear assemblies. Finally, all assemblies were screened for potential contamination using FCS-GX (Astashyn et al., 2024).

### Gene annotation

Gene annotation was obtained by the Ensembl team using their vertebrate genome annotation pipeline https://beta.ensembl.org/help/articles/vertebrate-genome-annotation.

### Repeat annotation

We used a modified version of pantera (Sierra and Durbin, 2024), panteraGA, that works in conjunction with FastGA (Myers et al., 2025) to obtain TE libraries from the combined 18 assemblies. As the TE libraries obtained with pantera will lack non polymorphic TEs, we added the TE families previously obtained following the protocol described in (Goubert et al., 2022), present already in Dfam (Storer et al., 2021). First we obtained an initial de novo library using RepeatModeler-2.0.3, with option-genomeSampleSizeMax 900000000 to sample the whole genome and obtain the most complete initial set of sequences (2,955 candidate TEs). We diverged from the protocol in that, instead of producing a priority list to then proceed to the manual curation of each candidate, we applied a de novo method to automatically extend the sequences looking for the whole sequence of the TE, described in the next section, branching the search if the initial seed appeared to belong to more than one family. The sequences obtained in this way were clustered with CD-HIT 4.8.1 (Li and Godzik, 2006) using parameters -aS 0.8 -c 0.8 -G 0 -g 1 -b 500 to define families according to the (Wicker et al., 2007) framework, sometimes known as the 80-80-80 framework. For the resulting consensus sequences, we checked for the presence of structural features, LTRs, TIRs and polyA sequences. We also obtained their ORFs and the products of them. We used BLAST 2.12.0 (Altschul et al., 1990) to compare the sequences to the existing RepBase annotations, to confirm which ones were already present. Finally, we used RepeatClassifier-2.0.3 (Flynn et al., 2020) to obtain an initial classification of the sequences, and proceeded to discard those for which their structural features did not indicate a complete element, in the case of families with well defined structures. We kept any large palindromes or elements with TIRs as putative non autonomous DNA transposable elements, but did not try to assign them to any particular family.

### Alignments, synteny, dotplots

The synteny plot was obtained using mumemto (Shivakumar and Langmead, 2025). Other inter-species alignments and self alignments were created with FastGA (Myers et al., 2025). Dotplots for sequences larger than 100kb were made with ModDotPlot (Sweeten et al., 2024). For smaller sequences our own R script was used to plot identity of k-mers size 13. A modified version of ModDotPlot was used for some plots so as not to include reverse complement matches in the identity.

### Annotation of tandem arrays and centroids

We identified putative centromeric regions as follows: using the identification of satellite sequences obtained with panteraGA, we selected the satellite sequence regions with the highest number of annotations on each chromosome as centromere candidates. We used TRASH (Wlodzimierz et al., 2023a) and FasTAN (https://github.com/thegenemyers/FASTAN) to confirm the boundaries of the centromere candidates and obtain the sequences of all satellites. For each chromosome in each species there was a single large region dominated by tandem satellite repeats of the same 237 bp repeat unit. We used FasTAN to obtain an accurate annotation of each tandem array. We used ModDotPlot (Sweeten et al., 2024) to obtain stained glass plots of the tandem arrays. We modified the code of ModDotPlot to generate different tandem arrays for each orientation of the satellites and confirm their distribution. We used our own R script to generate selfdotplots of the centroids. From TRASH we obtained the individual satellite sequences to produce satellite families.

### CpG methylation annotation

We used Oxford Nanopore’s dorado basecaller with options sup,5mCG_5hmCG to obtain DNA modified bases from the pod5 files. Used samtools to index and sort the reads and then modbam2bed with options -e -m 5mC --cpg to extract modified base counts. The results were plotted as averages of overlapping 20kb segments using an R script.

### Satellite clusters

For each centromere we extracted consecutive segments of 2370 bases, and obtained their 5-mer composition. Then we clustered the segments according to the presence and absence of the different 5-mers using the function otu from the kmer R package.

## Data availability

Sequencing data and assemblies are available in INSDC under BioProject number XXX. For the purposes of review the assemblies are available at https://zenodo.org/records/20933961

## Acknowledgements

We thank Richard Zatha, Sulla Manduwa, Heals Kabowa, Joel Elkin and Domino Joyce for assistance with sample collection, the Department of Fisheries of the Government of Malawi for facilitating Malawi cichlid specimen collection, Francesca Tricomi and Leanne Haggerty of the Ensembl team, EMBL European Bioinformatics Institute for gene annotation of the *Astatotilapia calliptera* genome, and Rob Martienssen and Felipe Karam-Teixeira for helpful comments. We acknowledge funding from the Wellcome Trust award 317408/Z/24/Z, and from the European Union’s Horizon 2020 research and innovation programme under the Marie Skłodowska-Curie grant agreement No 956229. For the purpose of open access, the author has applied a CC BY public copyright licence to any Author Accepted Manuscript version arising from this submission.

## Author contributions

P.S. carried out the analyses on finished genomes, C.Z. and S.W.L. assembled the genomes, B.F. extracted DNA and carried out the sequencing, M.B and M.N. oversaw sample collection, R.D. supervised the work. P.S. and R.D. wrote the manuscript.

