## Supplementary Figure for "Near-complete genomes for nine haplochromine cichlid fishes reveal a novel centromeric satellite structure organised around a pair of inverted elements"

### Supplementary figures.-

**Supp. Figs. 8 to 29.**- Self dotplots of 3 MB centered on the centromere of each chromosome for each of the nine species.

**Supp. Fig. 30.-** Satellite sequences from one chromosome aligned keeping their relative order in the centromere. The satellites belong, top down, to a region 5' of external satellites, 5' centroid, 5' internal satellites, 3' internal satellites, 3' centroid, 3' external satellites. The vertical arrows indicate examples of nucleotide variants which appear to be particular of just one of the segments.

**Supp. Fig. 8.-** Centromeres of chr 1

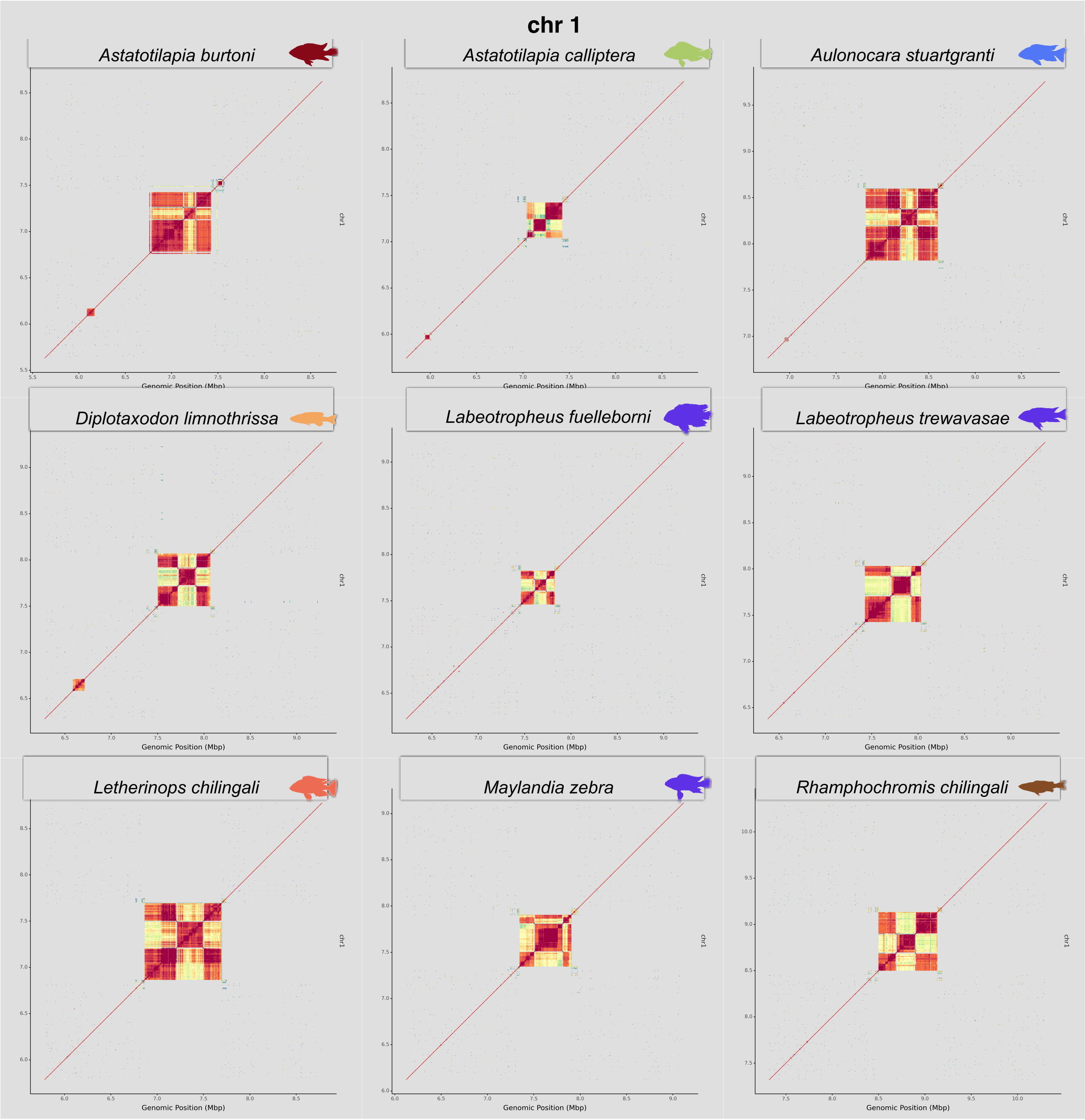

**Supp. Fig. 9.-** Centromeres of chr 2

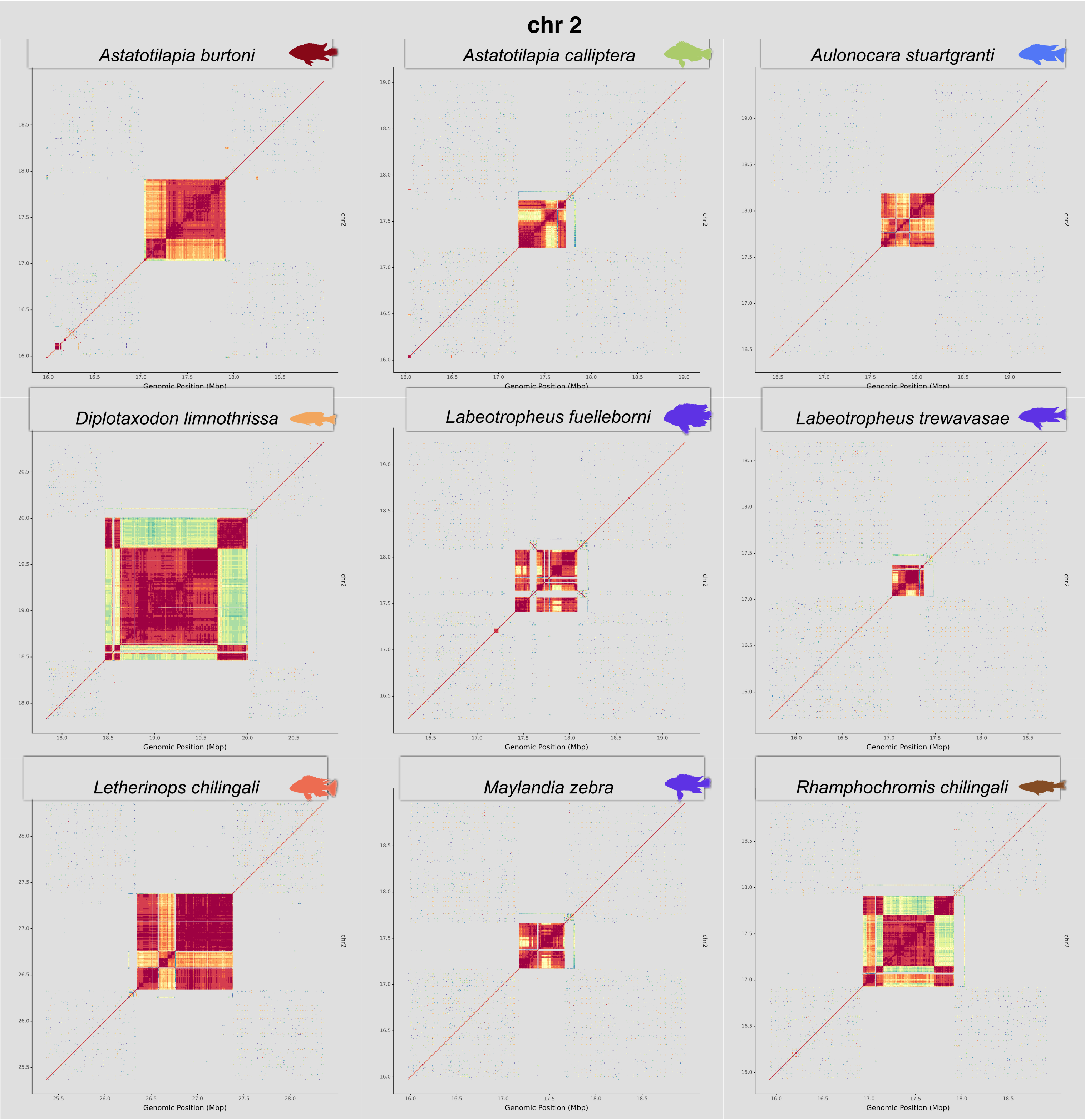

**Supp. Fig. 10.-** Centromeres of chr 3

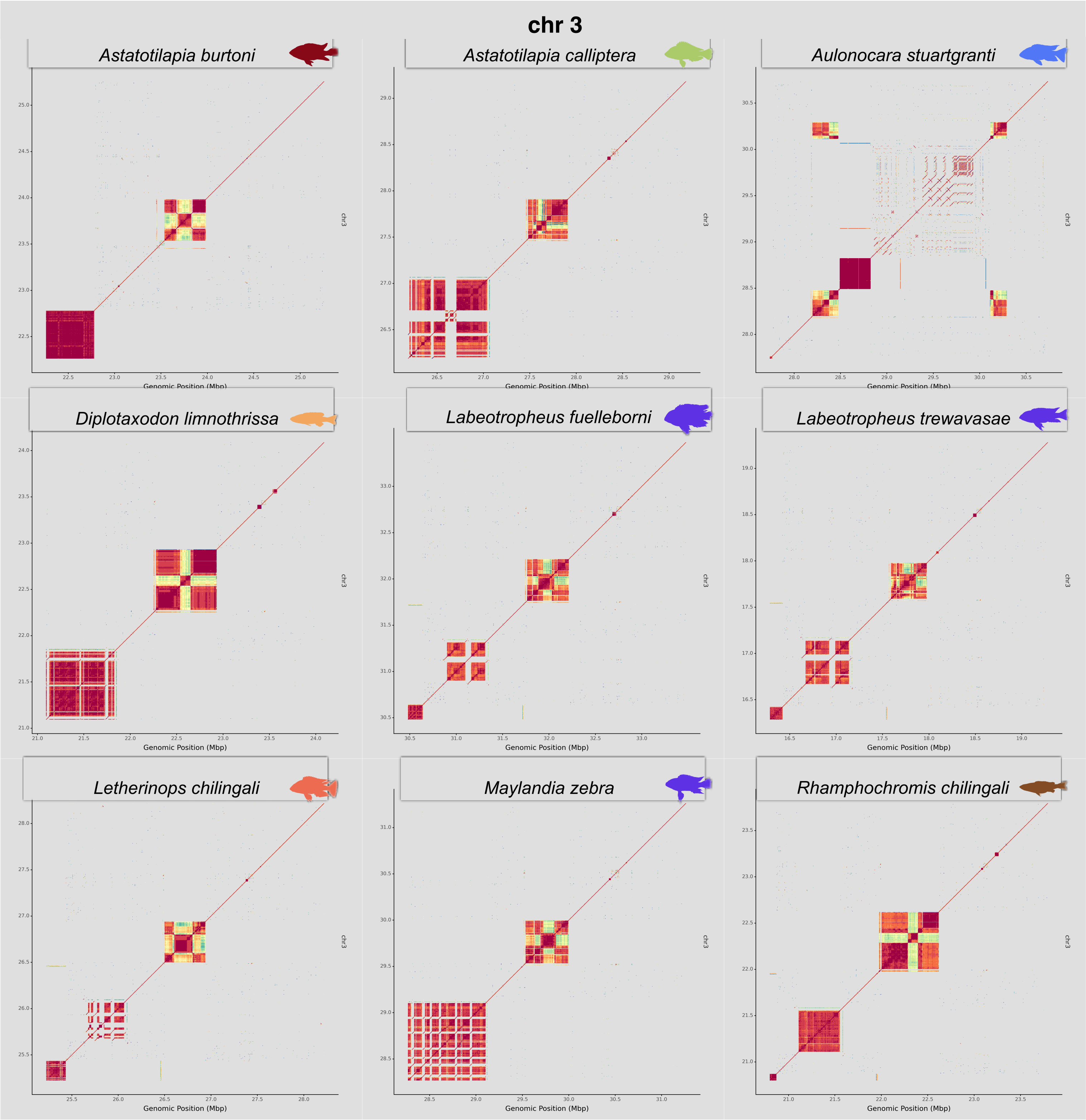

**Supp. Fig. 11.-** Centromeres of chr 4

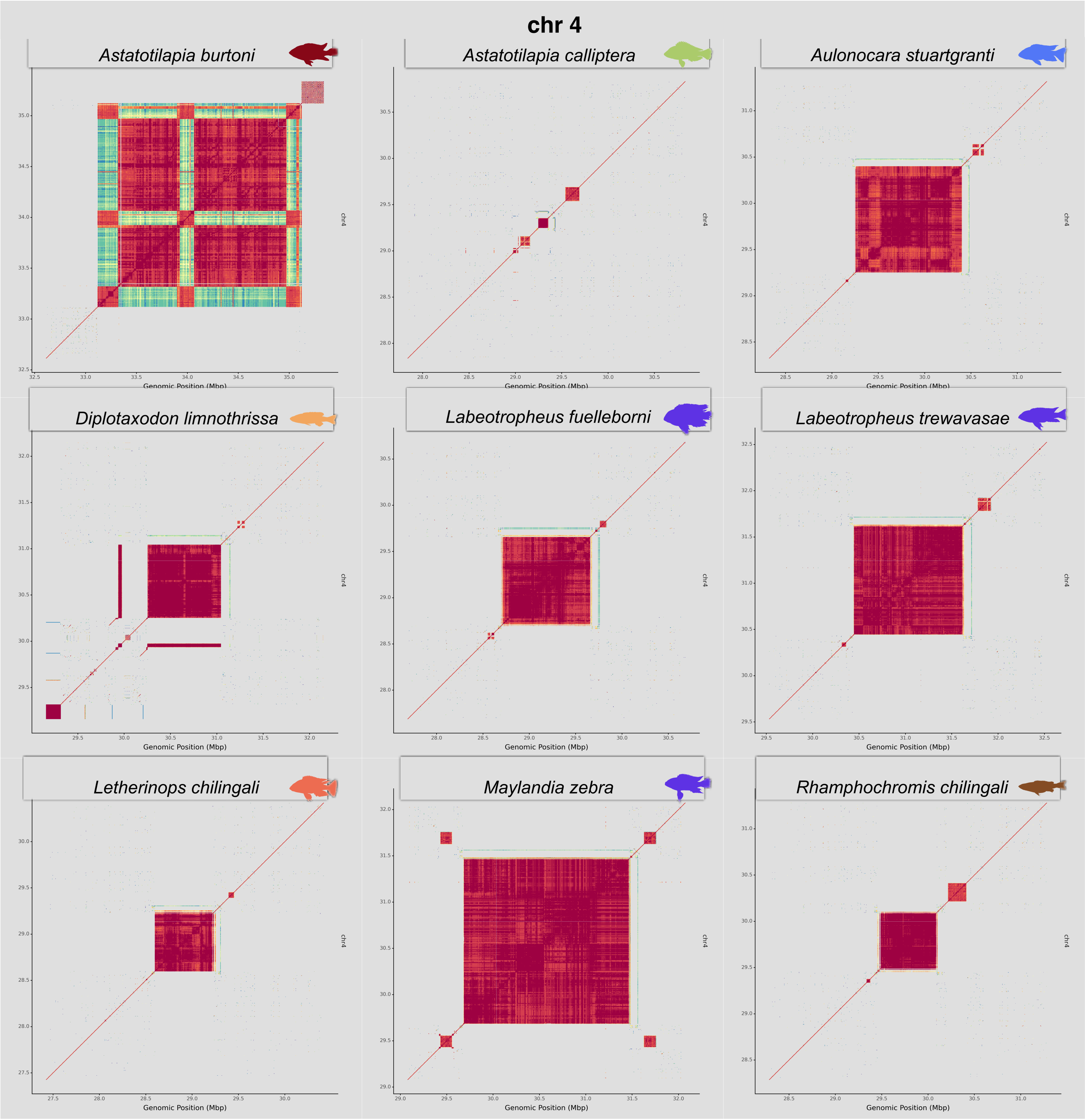

**Supp. Fig. 12.-** Centromeres of chr 5

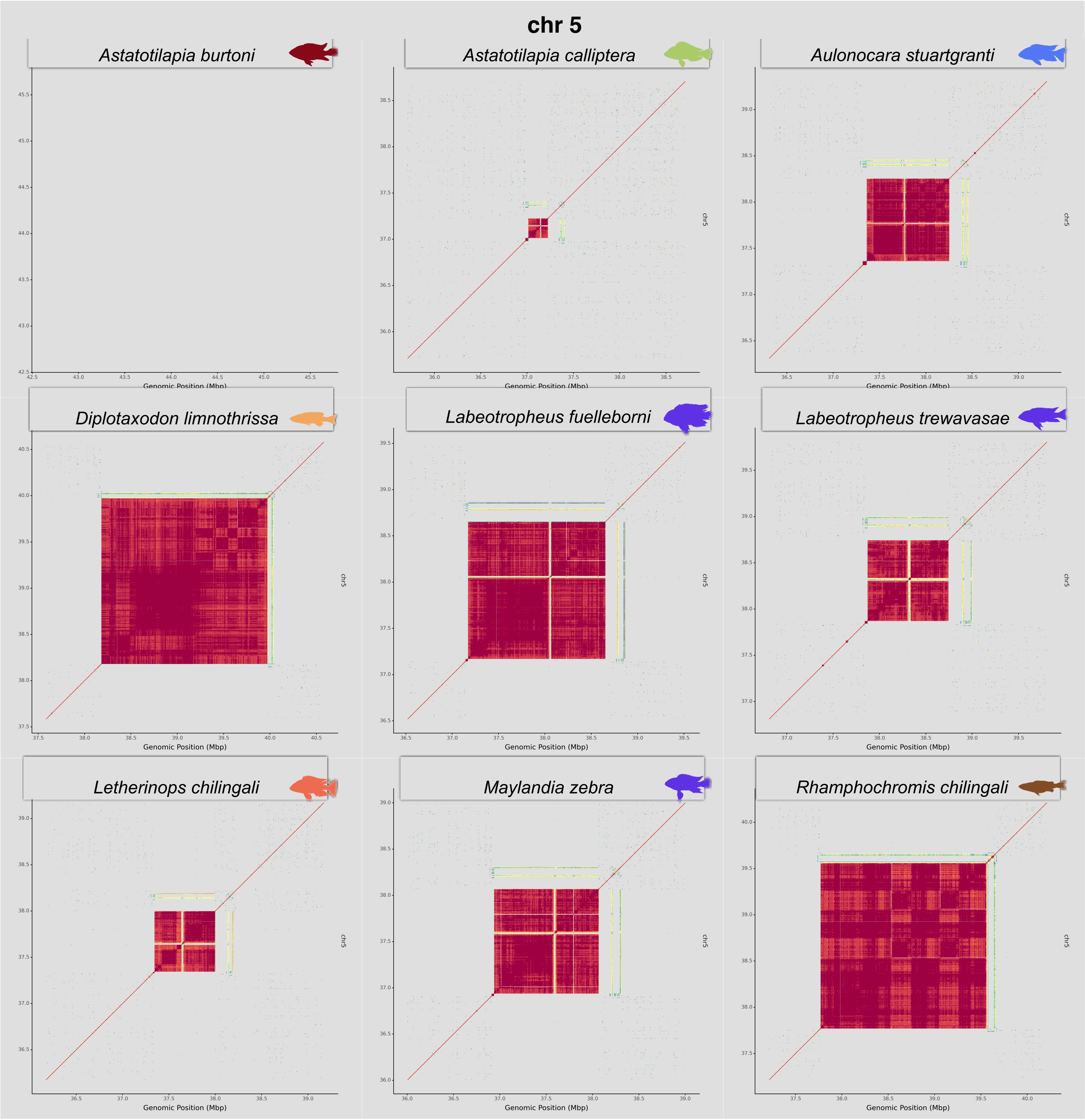

**Supp. Fig. 13.-** Centromeres of chr 6

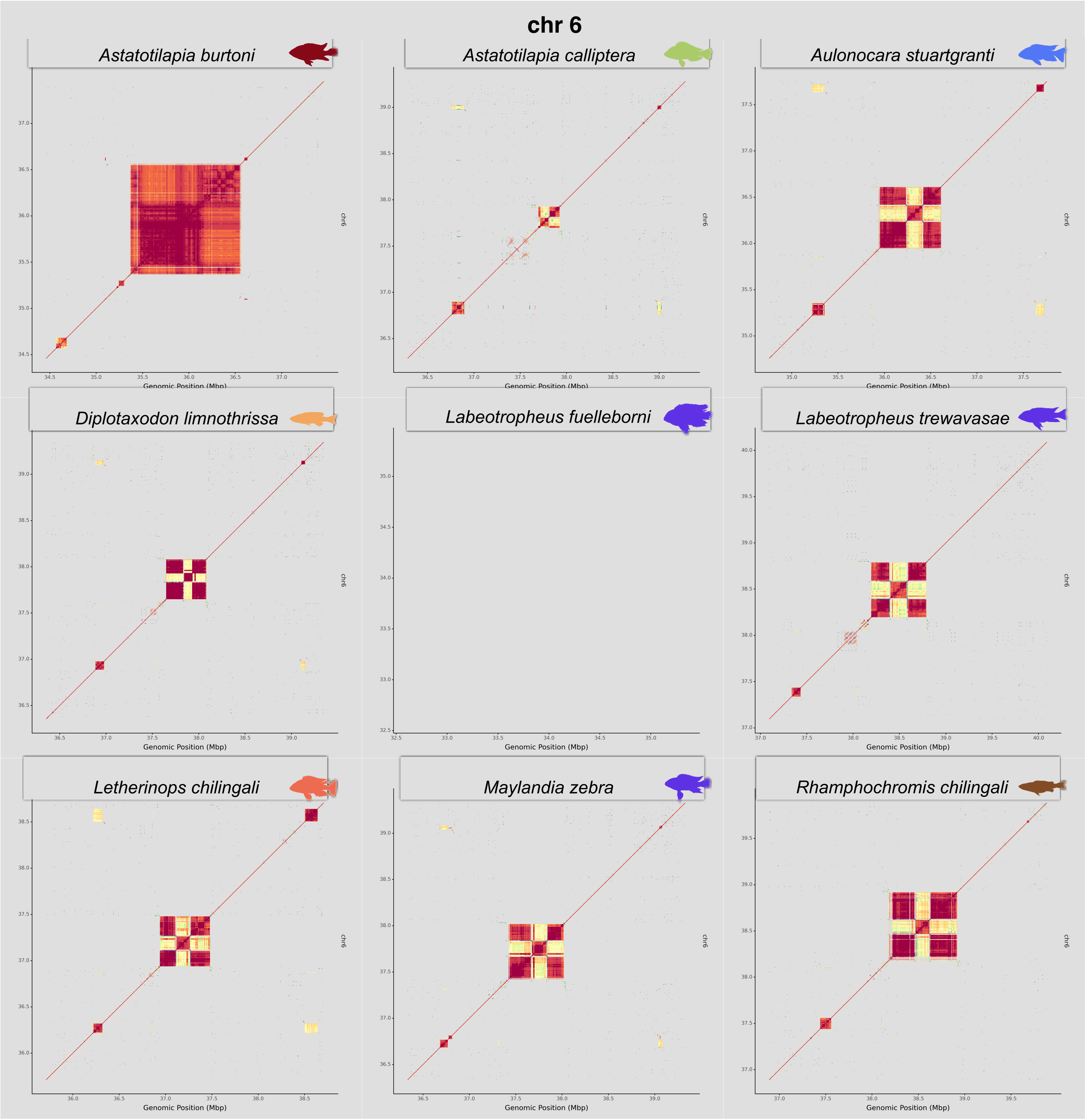

**Supp. Fig. 14.-** Centromeres of chr 7

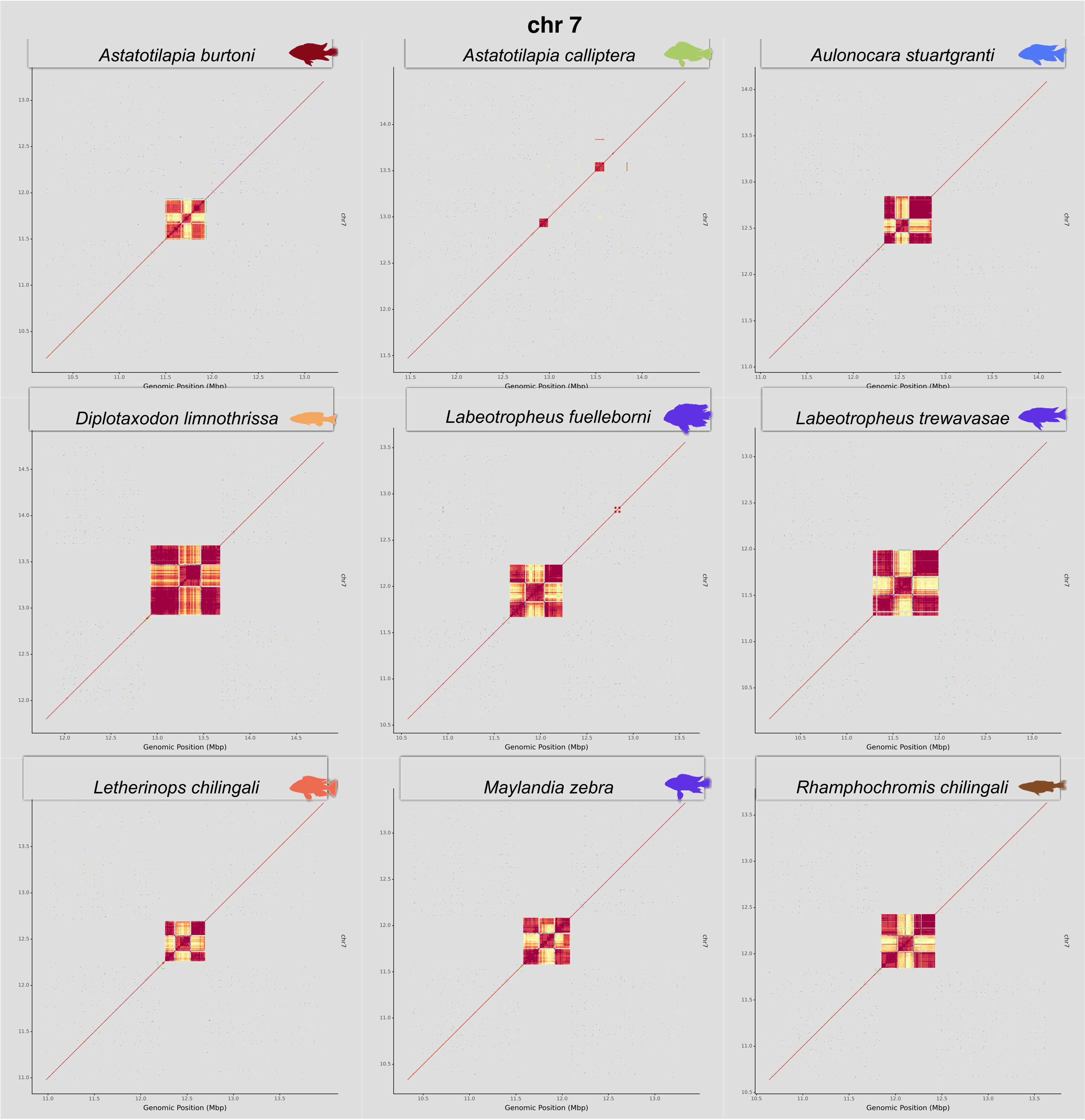

**Supp. Fig. 15.-** Centromeres of chr 8

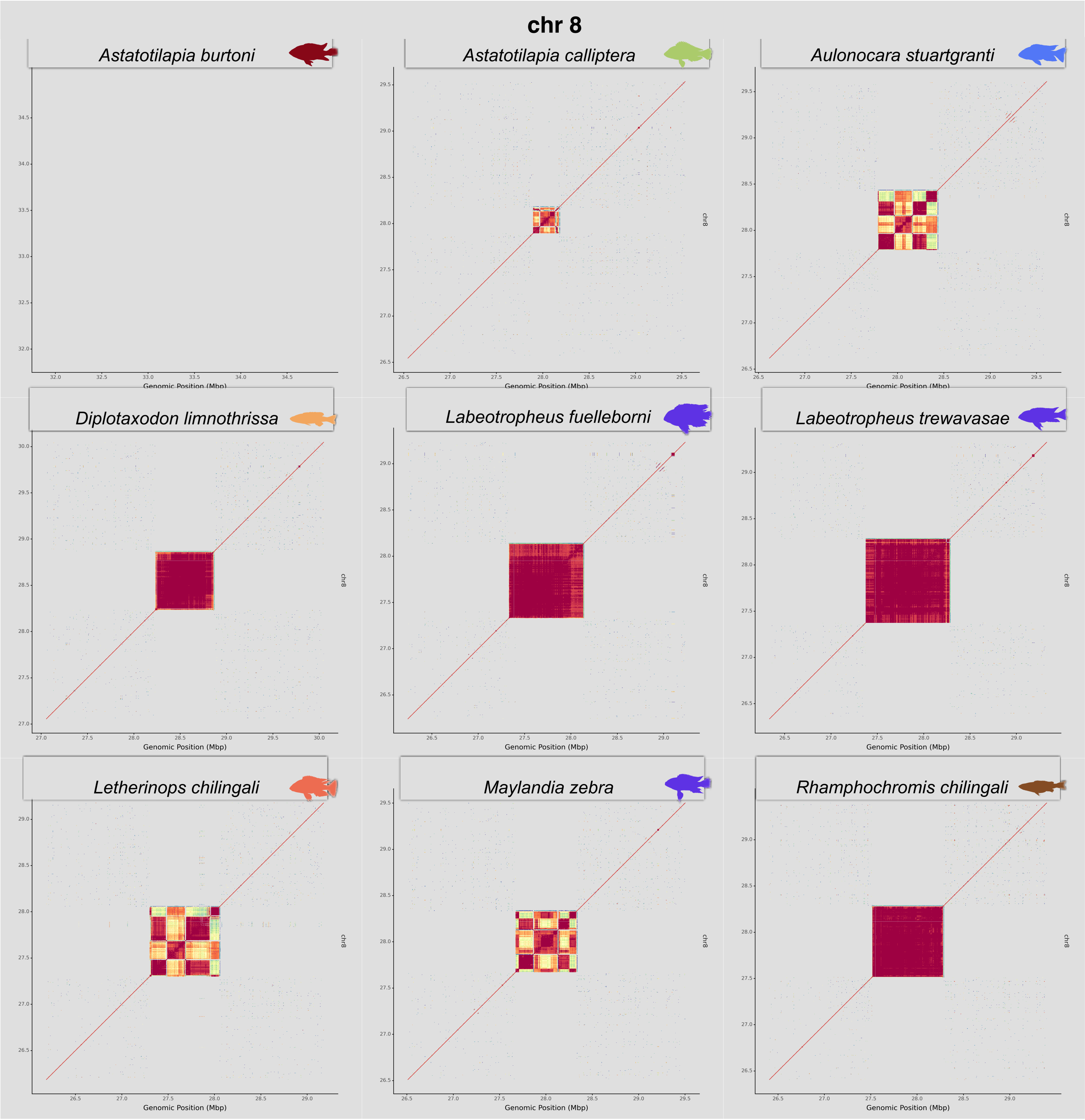

**Supp. Fig. 16.-** Centromeres of chr 9

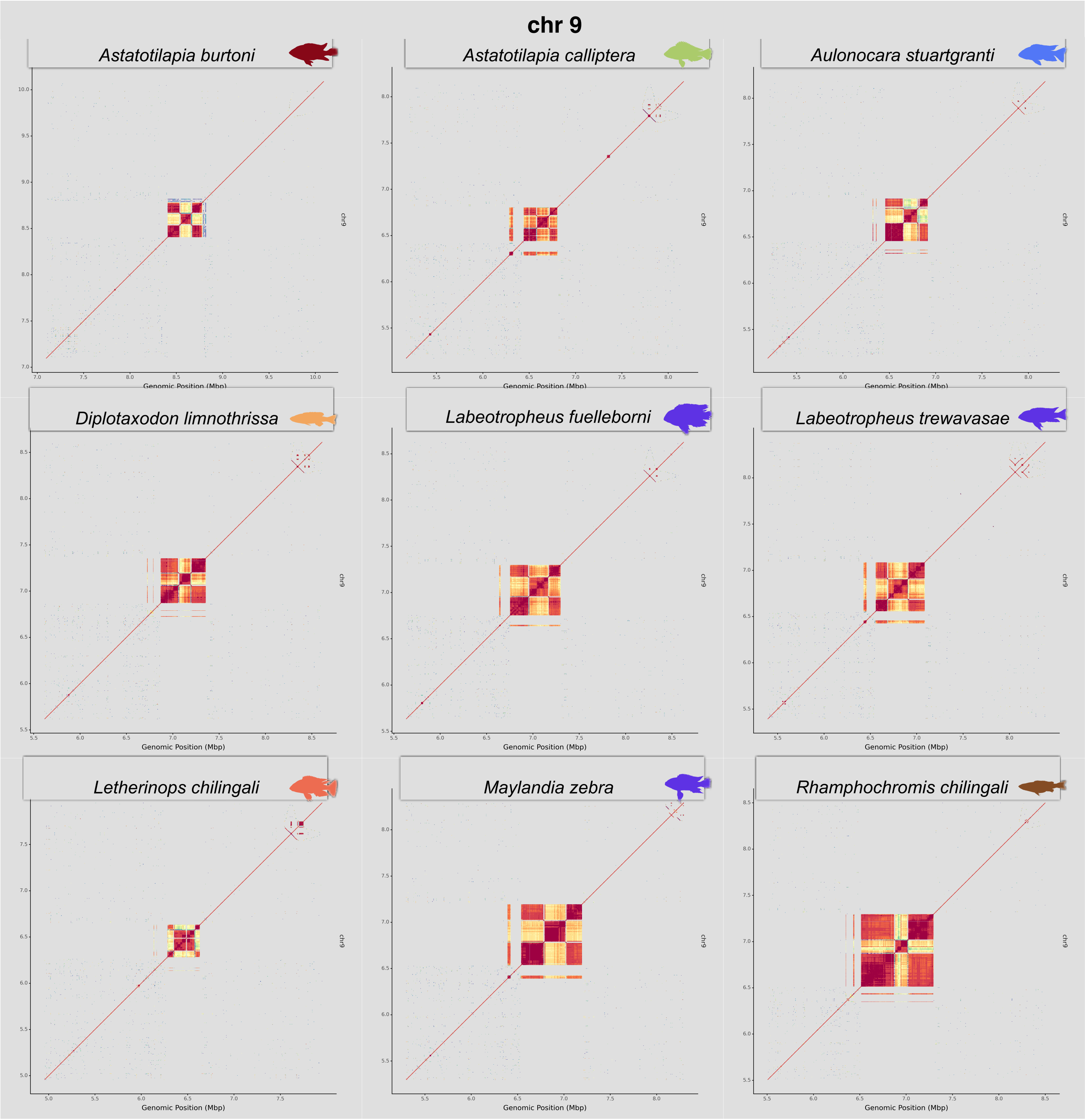

**Supp. Fig. 17.-** Centromeres of chr 10

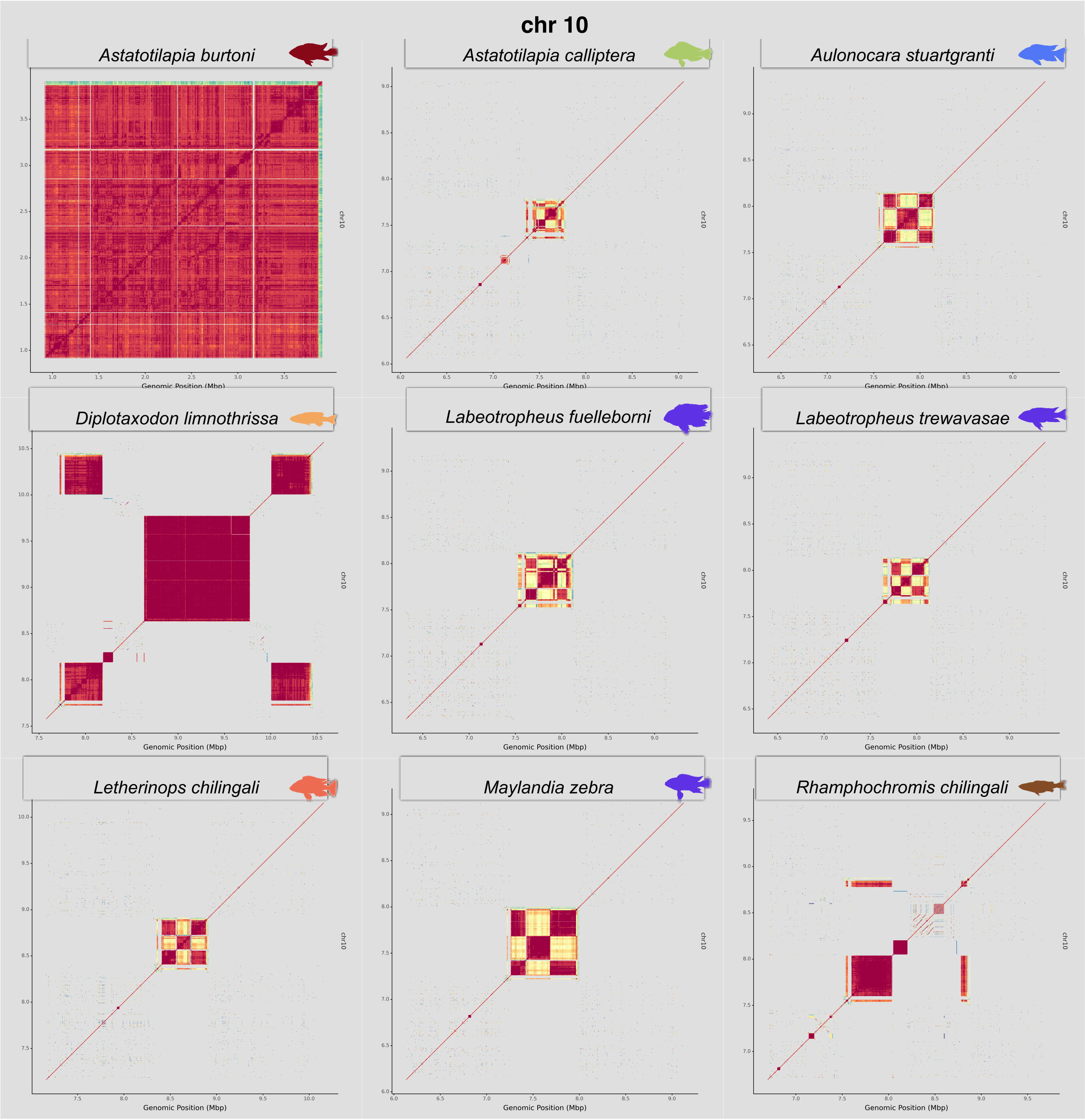

**Supp. Fig. 18.-** Centromeres of chr 11

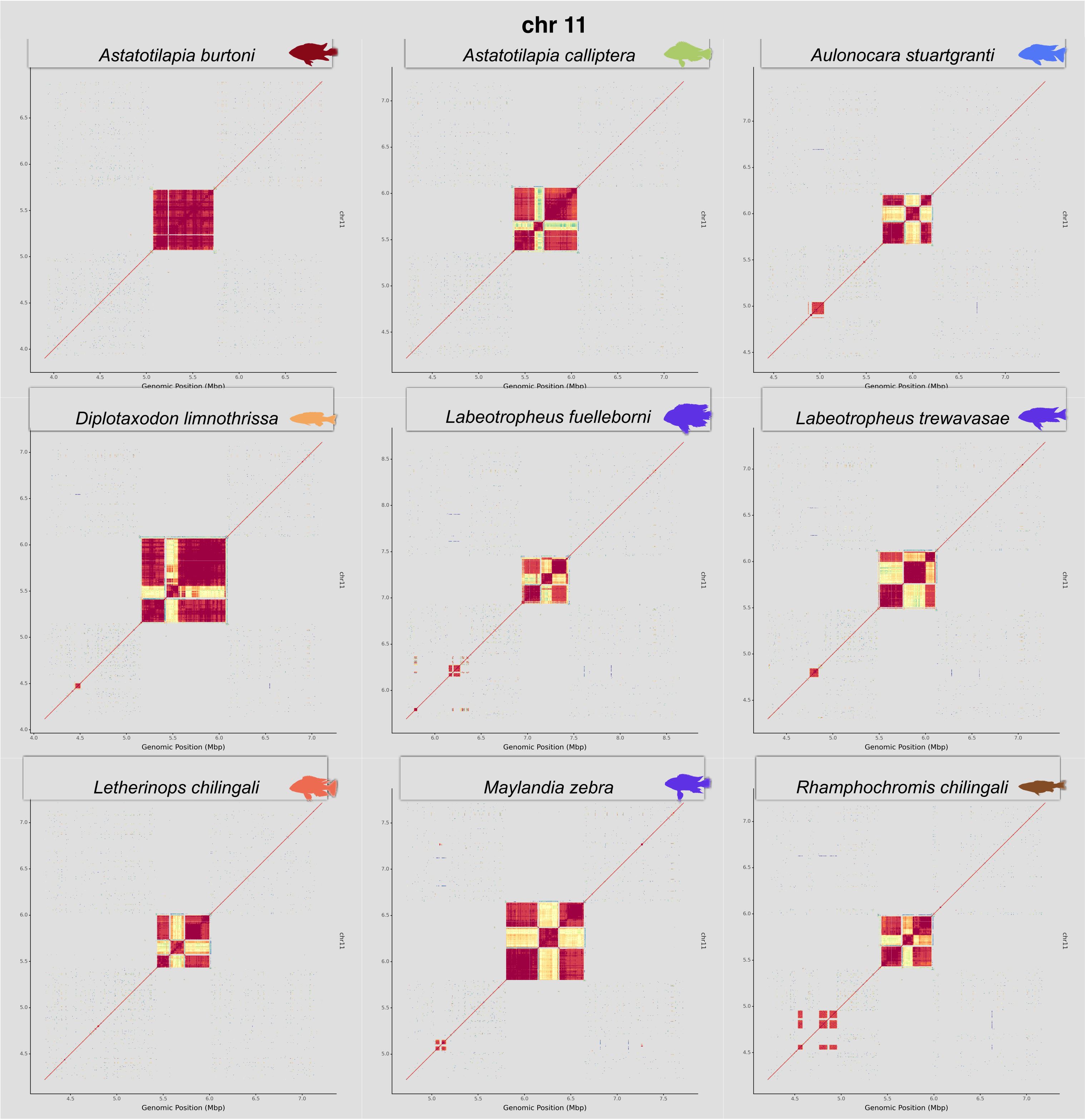

### **Supp. Fig. 19.-** Centromeres of chr 12

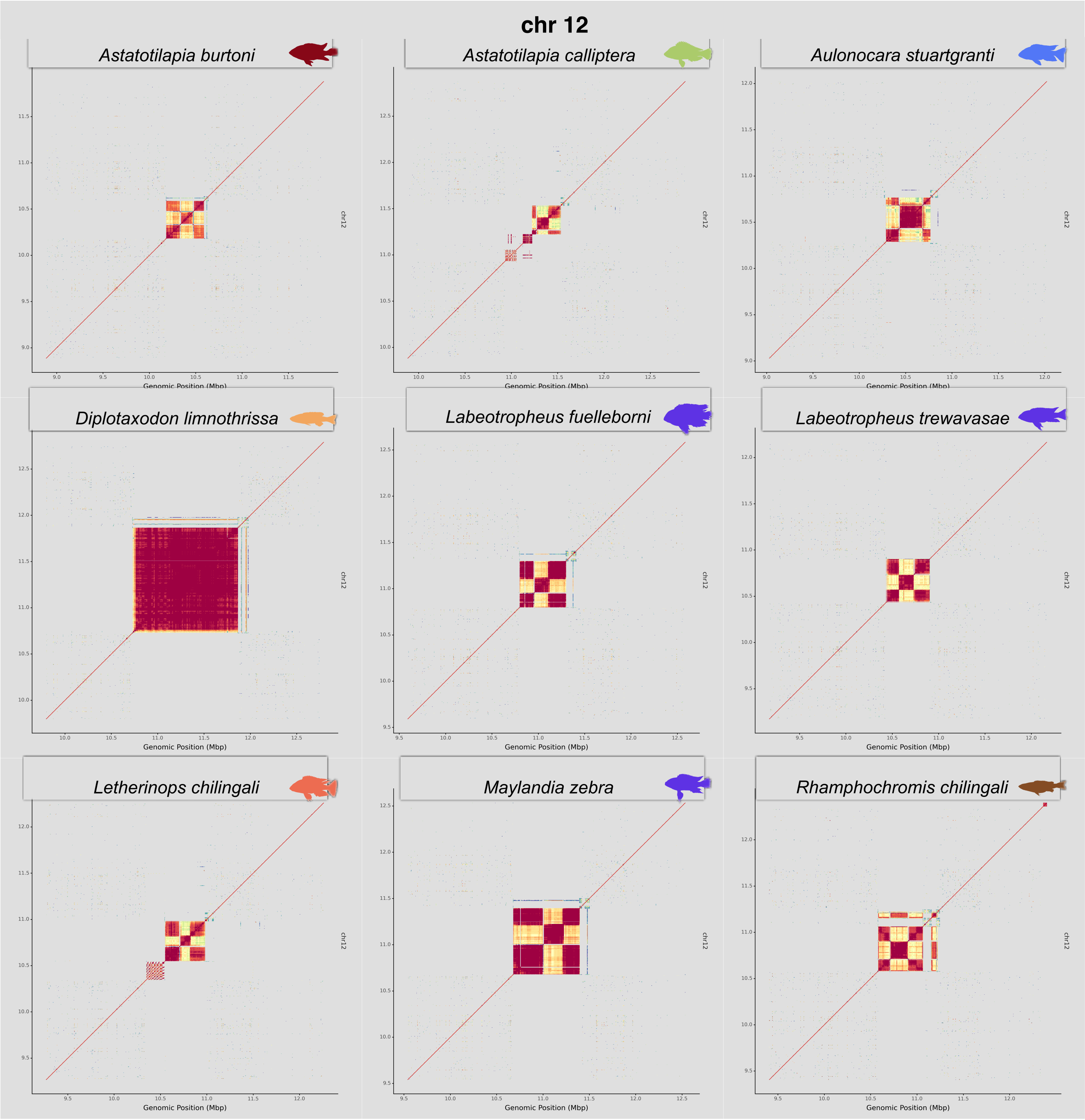

**Supp. Fig. 20.-** Centromeres of chr 13

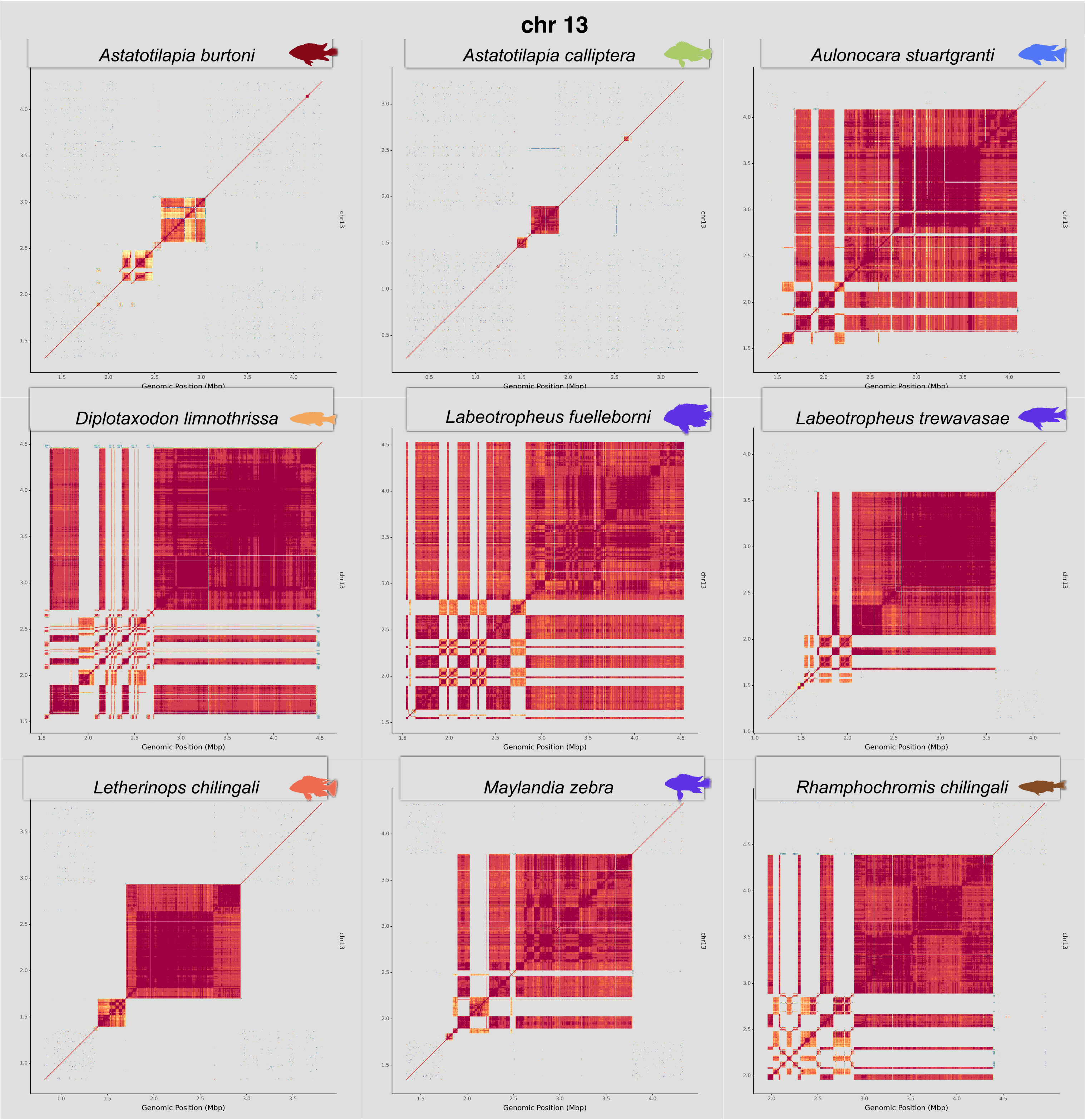

**Supp. Fig. 21.-** Centromeres of chr 14

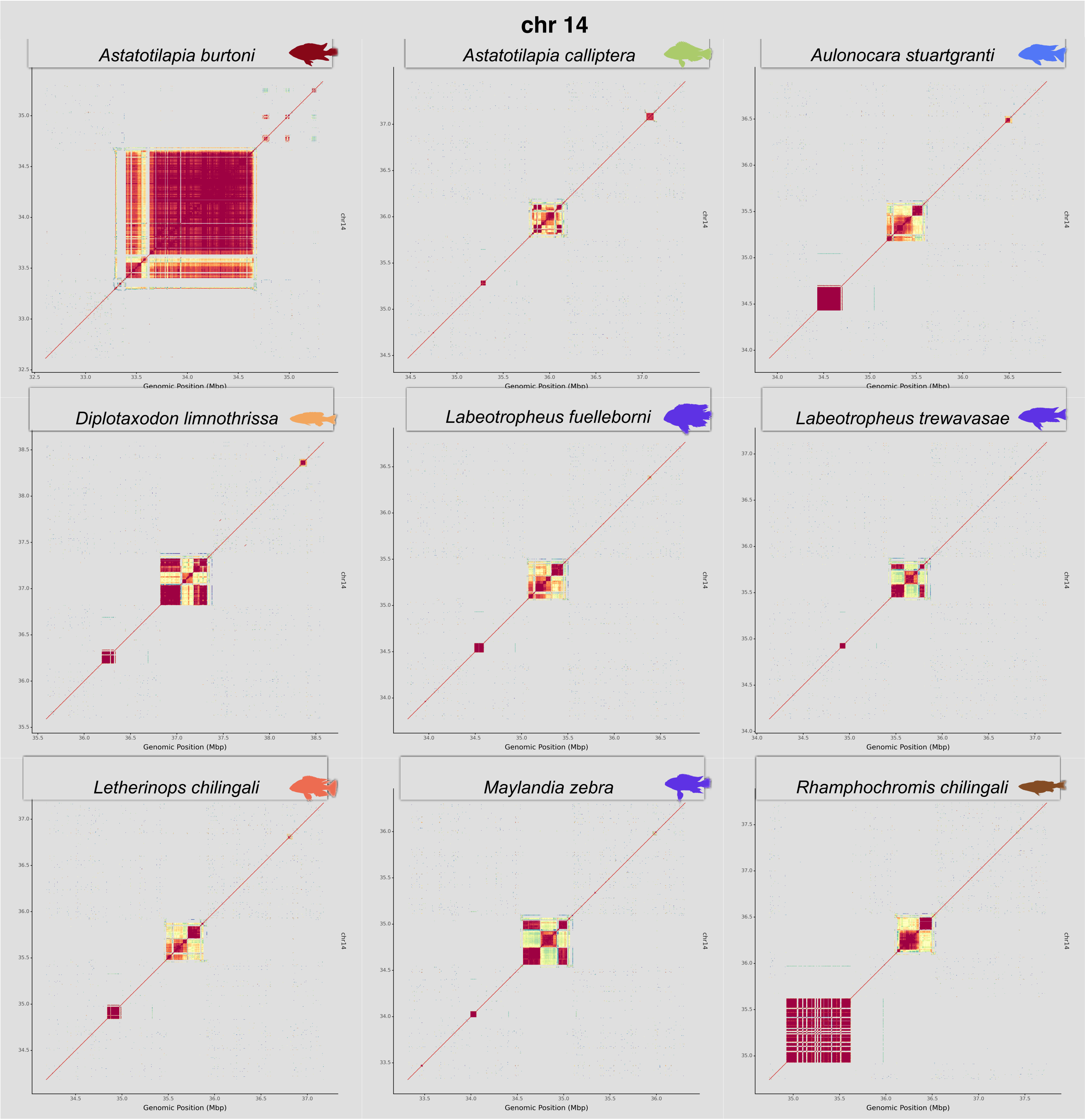

**Supp. Fig. 22.-** Centromeres of chr 15

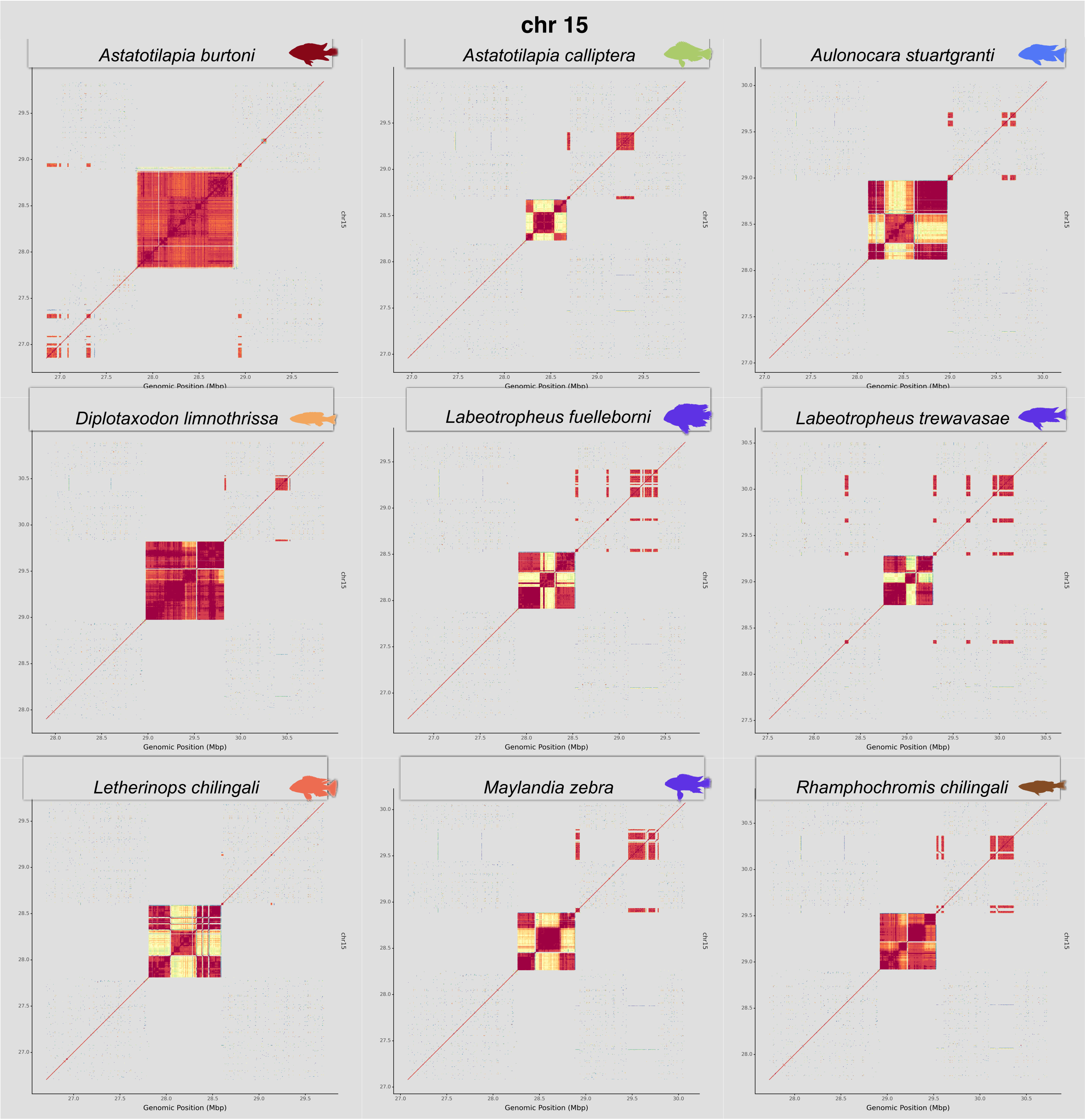

**Supp. Fig. 23.-** Centromeres of chr 16

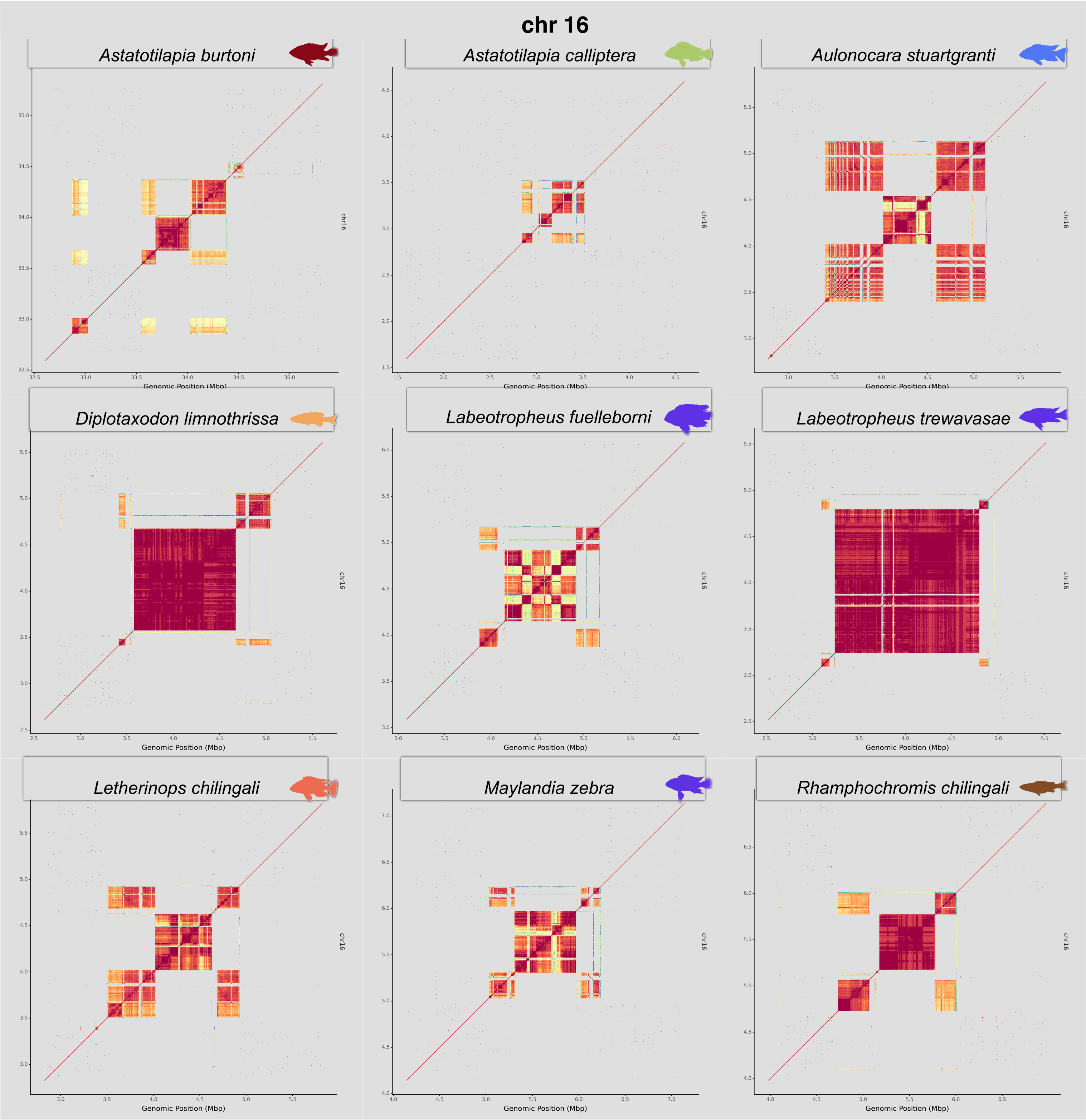

**Supp. Fig. 24.-** Centromeres of chr 17

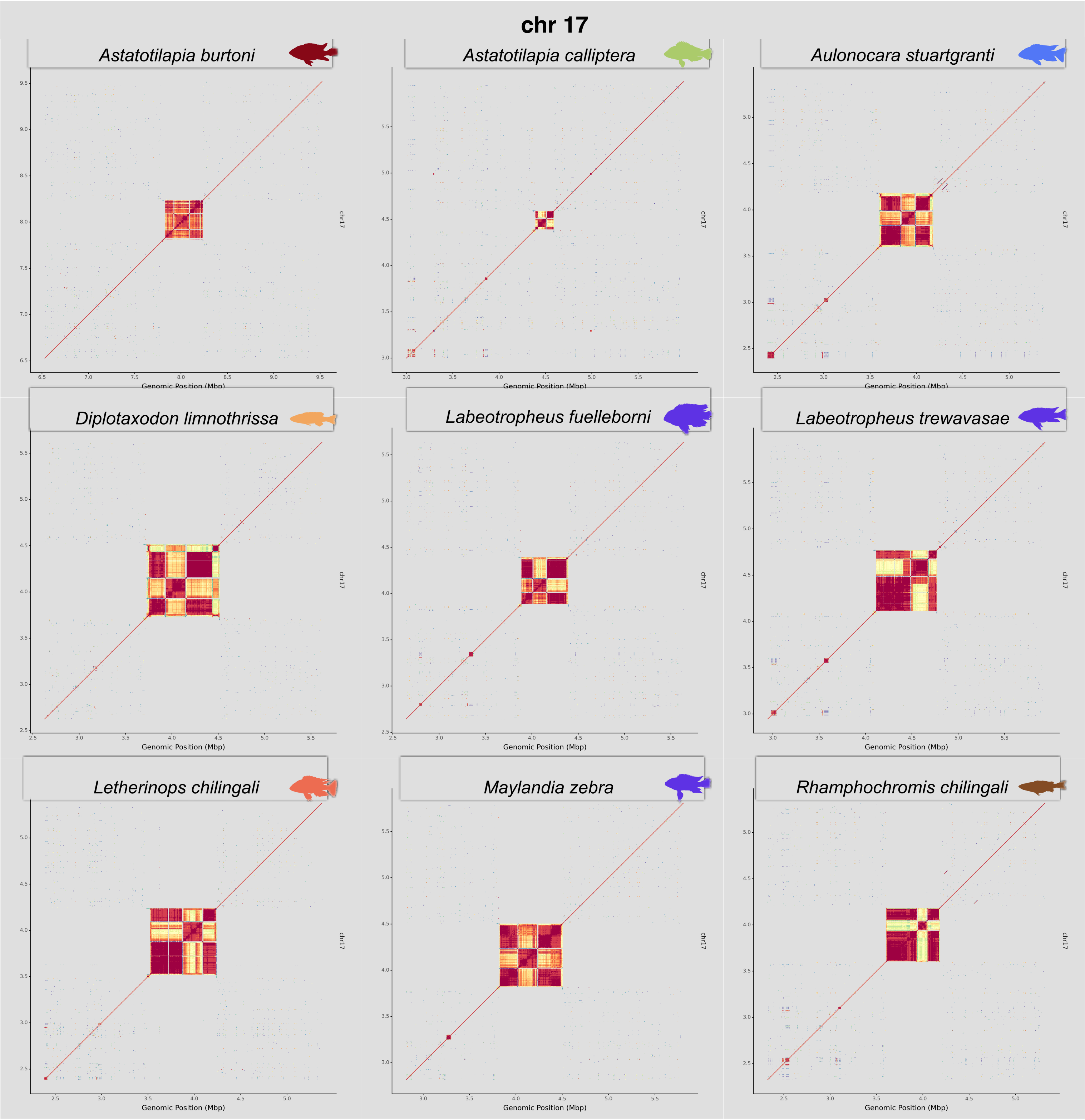

**Supp. Fig. 25.-** Centromeres of chr 18

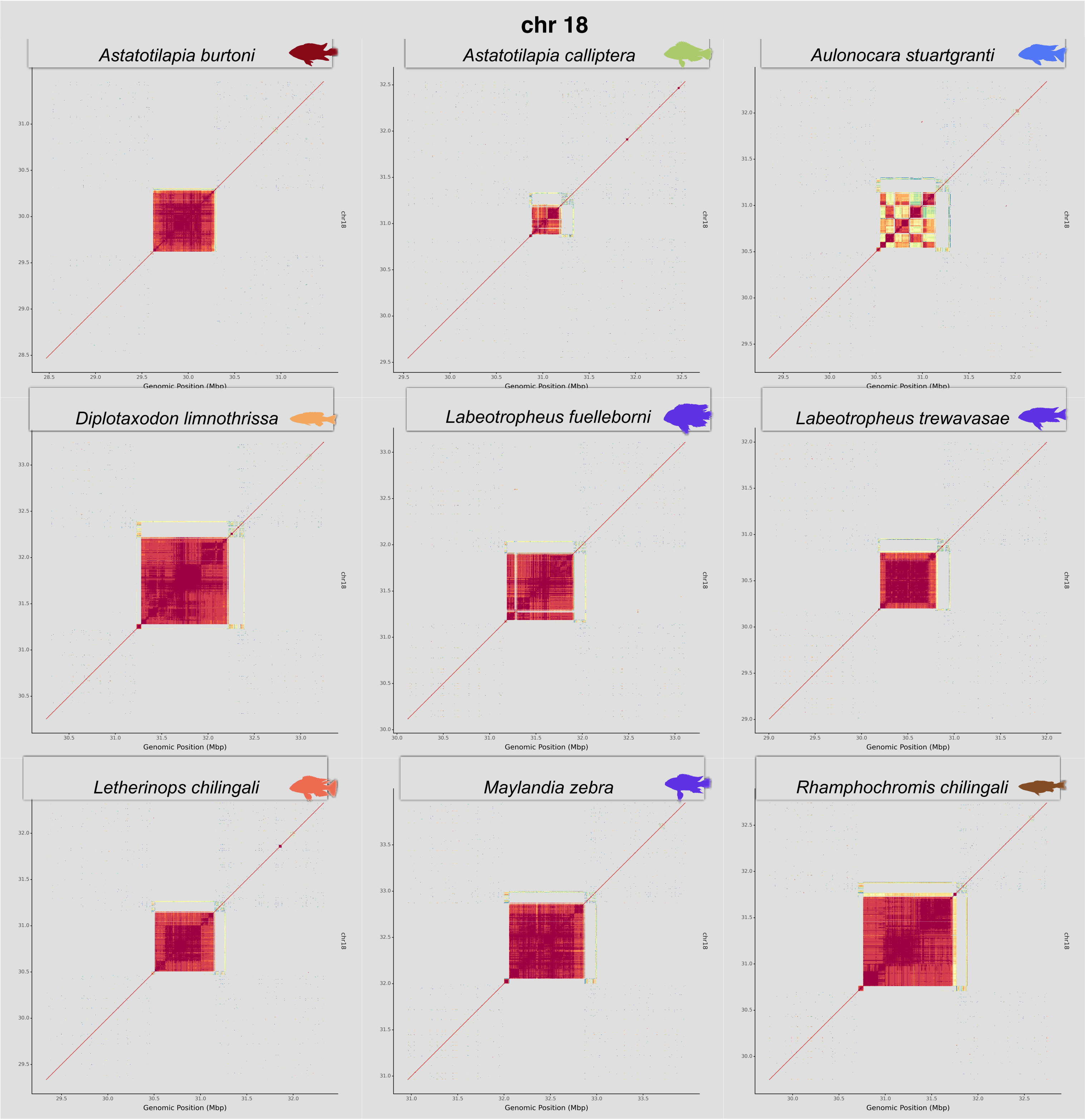

**Supp. Fig. 26.-** Centromeres of chr 19

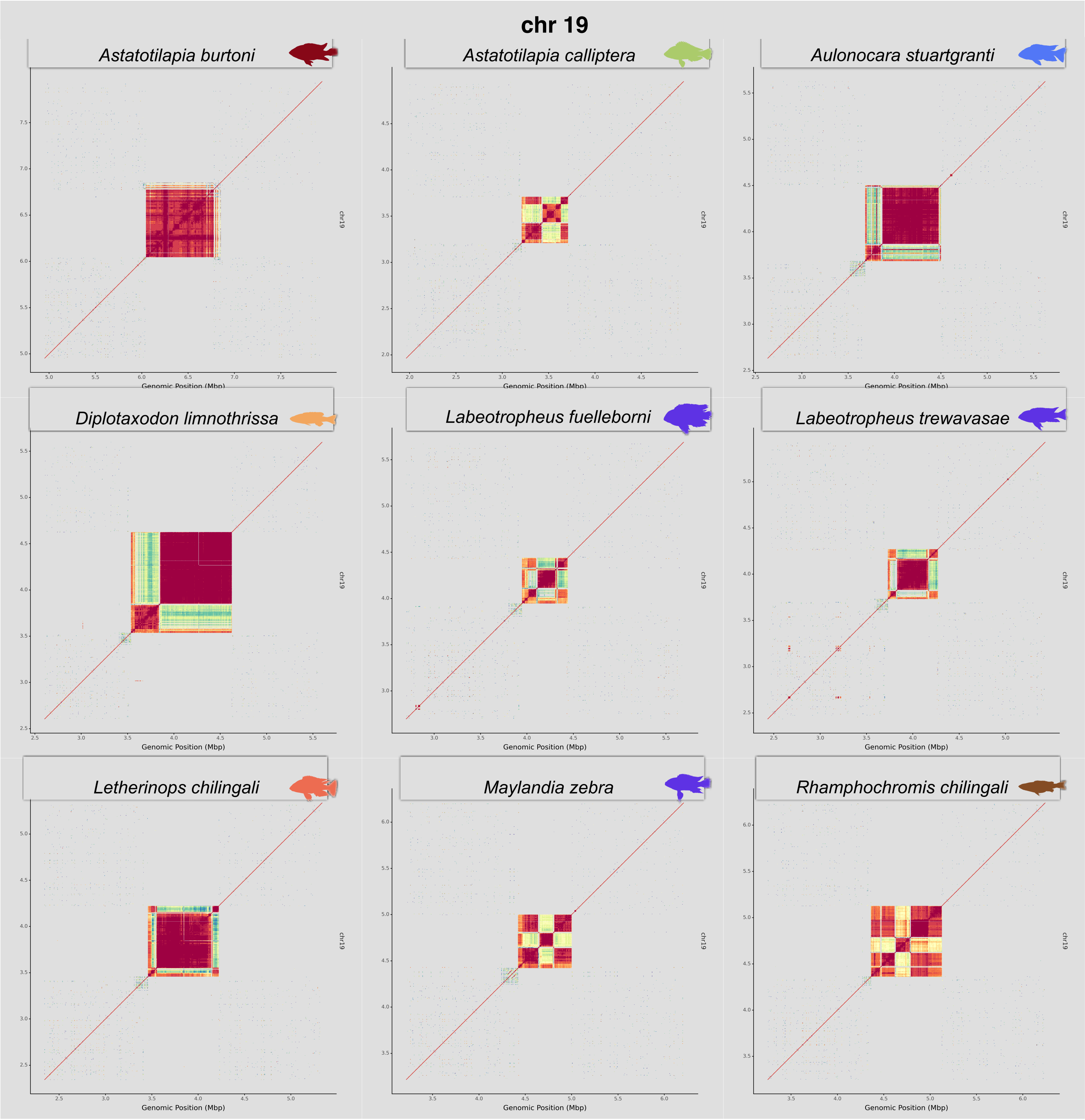

**Supp. Fig. 27.-** Centromeres of chr 20

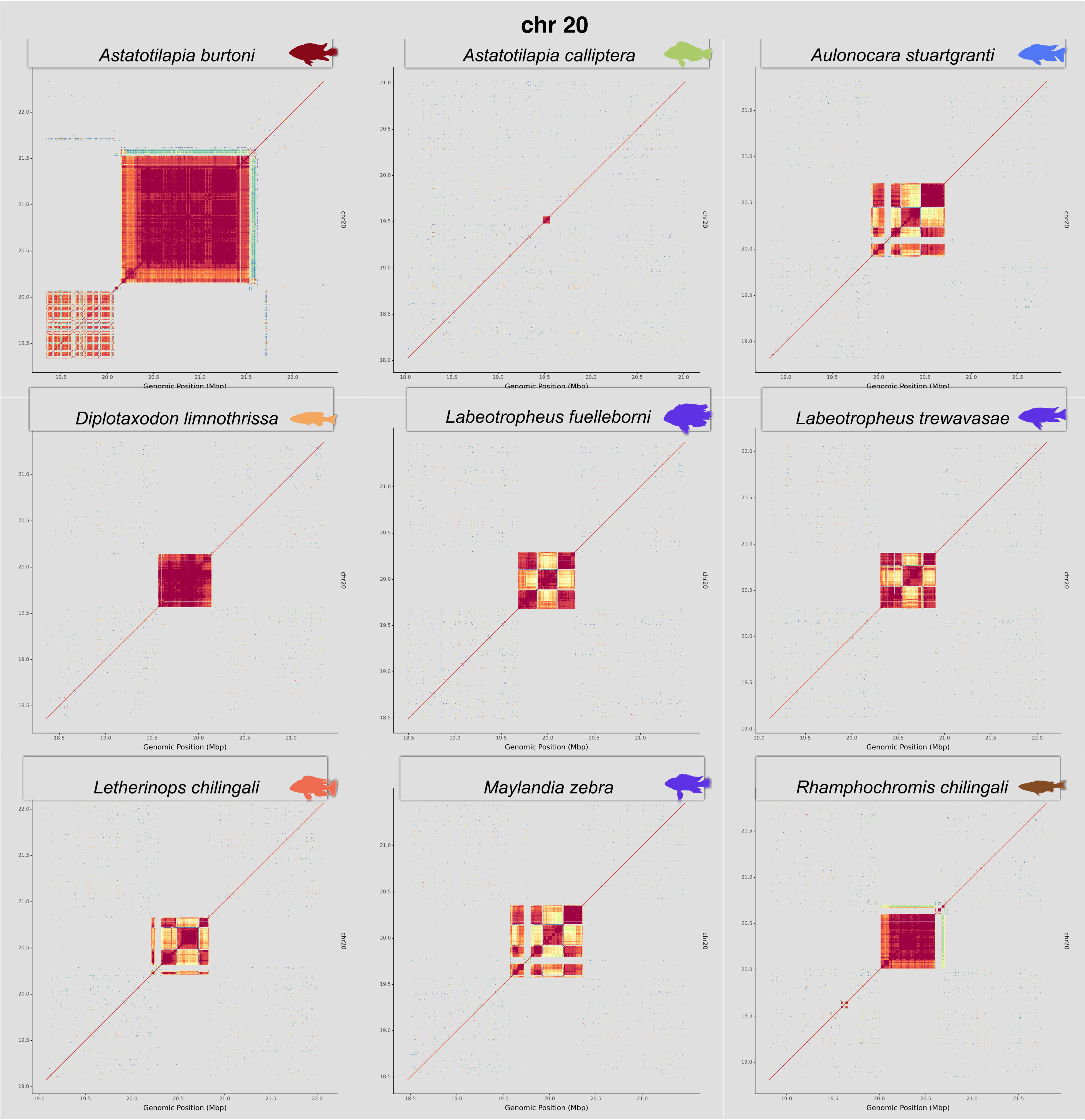

### **Supp. Fig. 28.-** Centromeres of chr 22

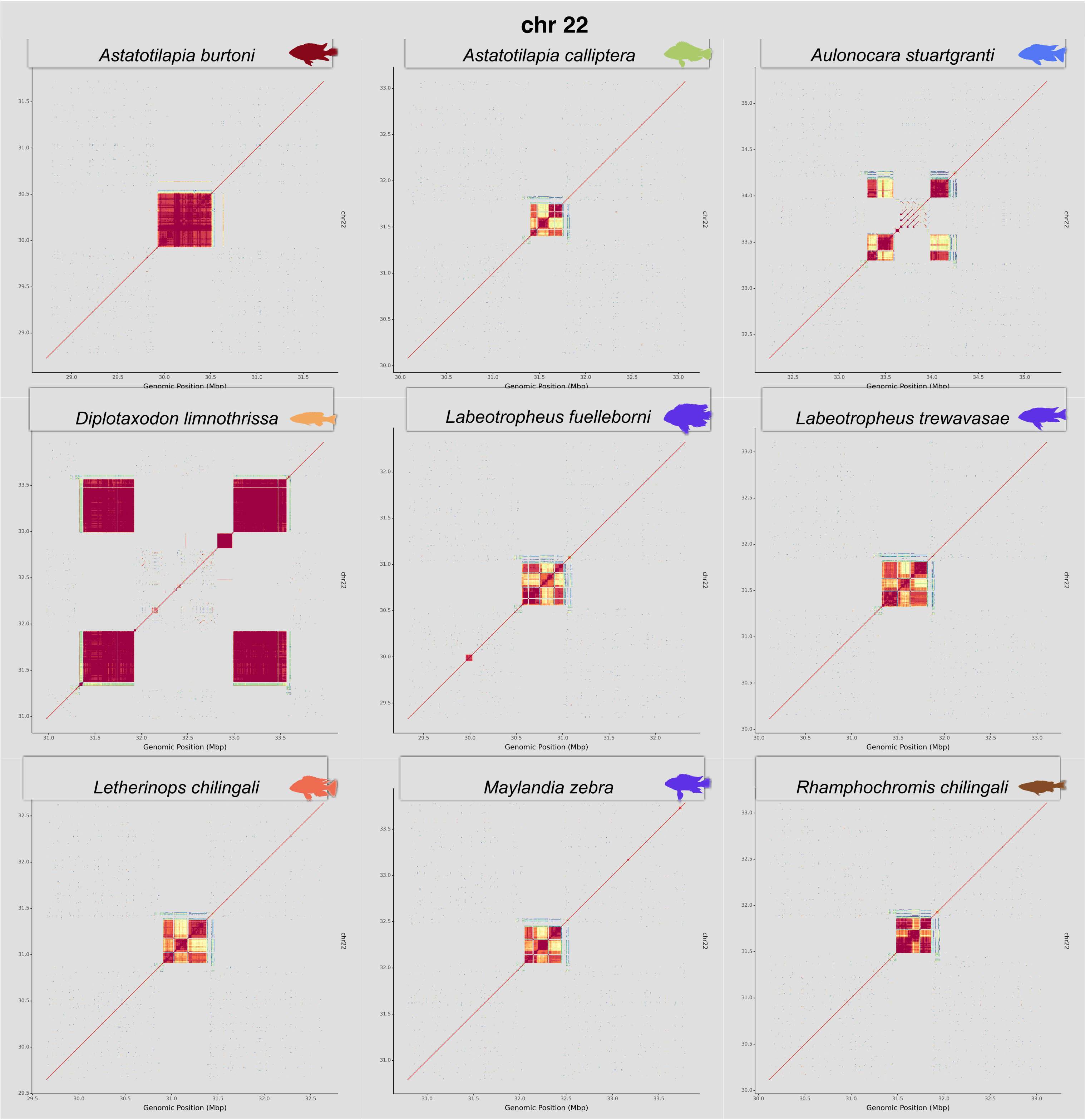

**Supp. Fig. 29.-** Centromeres of chr 23

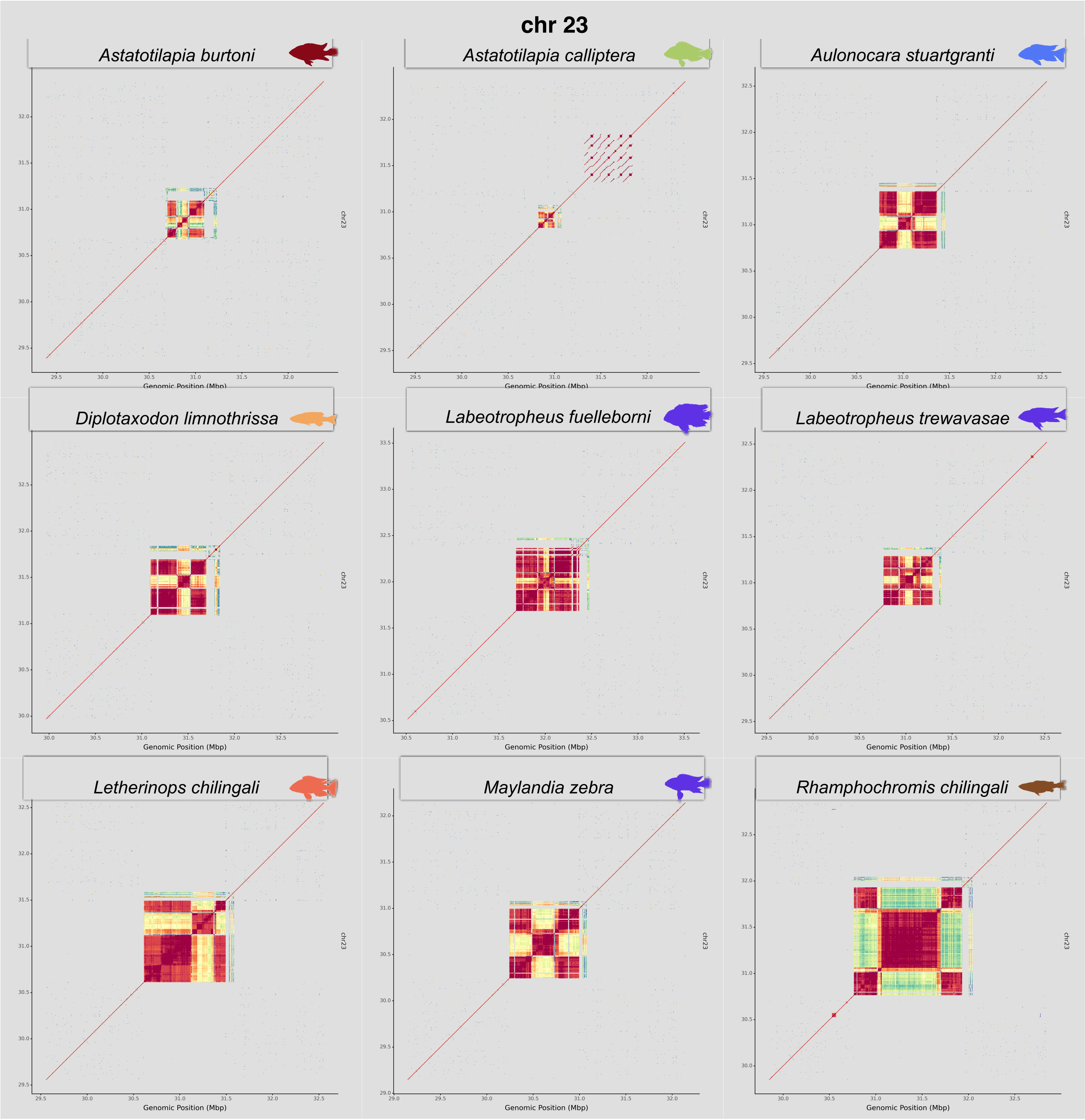

**Supp. Fig. 30.-** Cosegregating mutations in each of the four tandem repeats flanking centroids

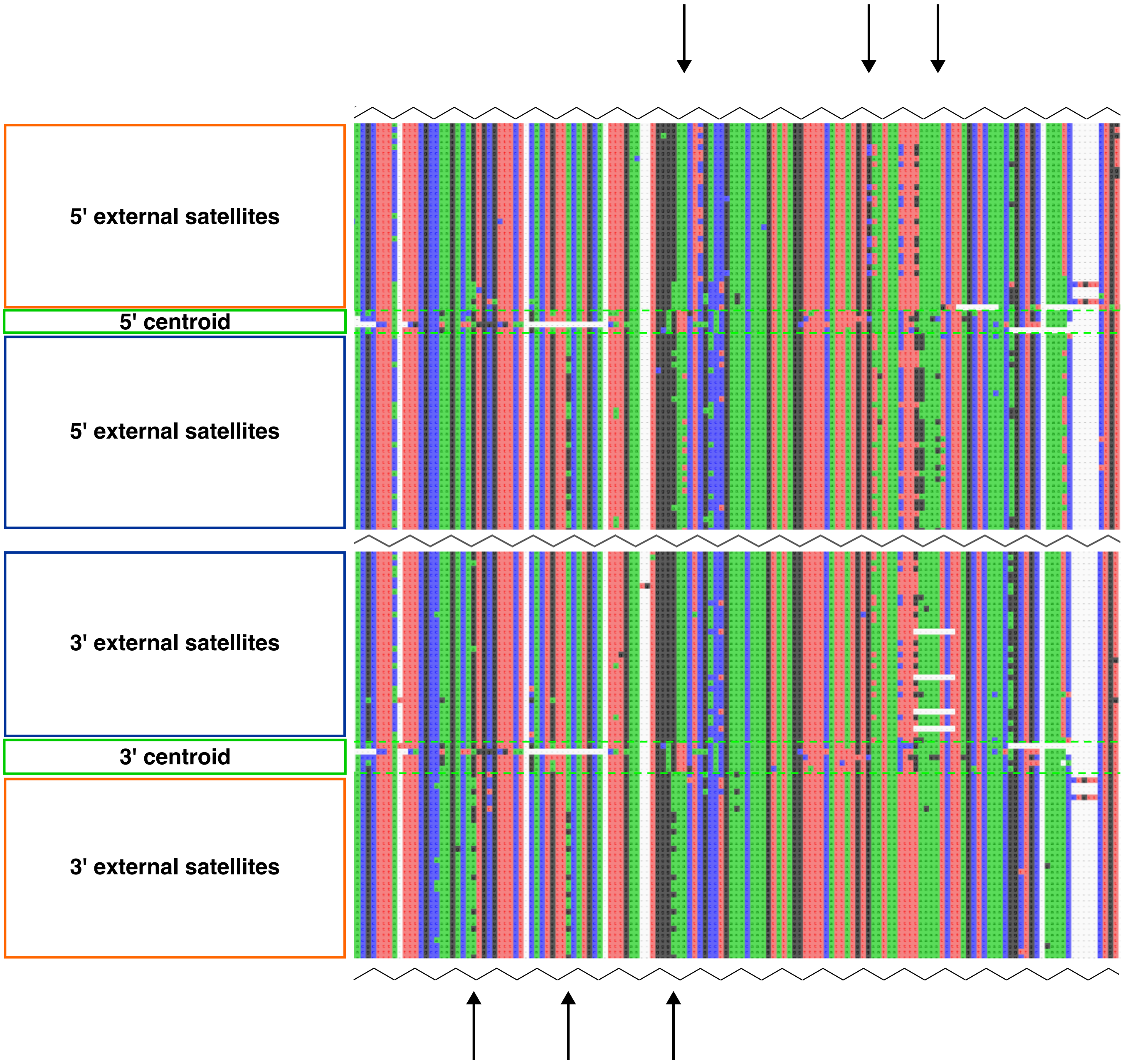
